# A critical contribution of cardiac myofibroblasts in right ventricular failure and the role of UCP2 SNPs in the predisposition to RV decompensation in pulmonary arterial hypertension

**DOI:** 10.1101/2025.07.25.666908

**Authors:** Yongneng Zhang, Sébastien Bonnet, Steeve Provencher, Alois Haromy, Yongsheng Liu, Yuan-Yuan Zhao, Sandra Breuils-Bonnet, Dawn E. Bowles, Michelle Mendiola Pla, Gopinath Sutendra, Evangelos D. Michelakis

**Author notes:** Correspondence to: Evangelos D. Michelakis, MD, Department of Medicine, University of Alberta, Edmonton, Alberta, Canada T6G 2B7.

## Abstract

The mechanism of transition from compensated (cRV) to decompensated right ventricle (dRV) in pulmonary arterial hypertension (PAH) is unknown. We explored the role of RV cardiac myofibroblasts (cMFB) on this transition utilizing a rat model and 3 cohorts of 81 patients which included clinical data, RV tissues and blood. We hypothesized that the loss of UCP2, critical for mitochondrial calcium (mCa^++^) regulation and cardiac fibroblasts (cFB) differentiation, is associated with dRV in rats and humans; and that a loss-of-function UCP2 SNP (rs659366) may predict dRV in human PAH. We separated rat cRV from dRV based on catheterization and echocardiographic criteria and found a significant increase in cMFB in dRV. In isolated hearts, RV contractility was lower in dRV but not in isolated cardiomyocyte (CM), pointing to a non-CM cause. Mitochondrial respiration was lower in dRV cMFB than in control and cRV cFB. mCa^++^ was progressively decreased from normal to cRV to dRV c(M)FB, and the same was true for c(M)FB (but not CM) UCP2 levels. Human PAH, but not secondary pulmonary hypertension, dRVs had more cMFB and less UCP2 than control and cRVs. Decreased UCP2 (protein and mRNA) levels and the presence of heterozygous/homozygous UCP2 SNP were associated with worse RV performance (TAPSE, cardiac index), even among patients with similar mean pulmonary arterial pressure. Our data point to a change of cell identity (cFB to cMFB) in the RV as a driver of RV decompensation. UCP2 SNPs are promising biomarkers for early cRV transition to dRV in PAH.

## Introduction

The main driver of mortality and morbidity in Pulmonary Arterial Hypertension (PAH) is the function of the right ventricle (RV) rather than the degree of rise in mean Pulmonary Artery Pressure (mPAP)^1,2^. In PAH, the compensating RV stage (cRV) is quite short, with decompensation occurring often in months^3^, compared to the left ventricular hypertrophy where the compensation can last decades^4^. Furthermore, between 2 patients with the same age/sex and mPAP, one may have a shorter cRV stage and decompensate faster than the other^5,6^, complicating PAH clinical management and timing of referral for transplantation. What makes the transition from cRV to decompensated RV (dRV) and why one RV more prone to failure than another is unknown. The presumption so far is that there is an energetic failure of cardiomyocytes combined with inadequate angiogenesis in the hypertrophied RV (i.e., relative ischemia), resulting in cell death, collagen deposition and structural remodeling^7,8^. In the left ventricle (LV), cardiac myofibroblasts (cMFB) increase quickly post myocardial infarction, mostly by differentiation from cardiac fibroblasts (cFB) in order to limit the myocardial wound via secretion of collagen and scar formation^9^. In contrast to cFB, cMFB express high levels of alpha-smooth muscle actin (α-SMA) that allows them to contract and migrate to the wound site, but also increase the stiffness of the ventricle and compromise its function^10^. cMFB can reach >30-40% of the cell population in the LV and also increase in other cardiac pathologies as well, like hypertrophy or diabetes^11,12^. There is evidence that differentiation and proliferation of cMFB can both be halted and even reversed^13^, providing a rationale for therapeutic targeting of cMFB. Although this has not been tested in the RV, it is very important because once dRV ensues, any improvement in the pulmonary vascular remodeling and mPAP through PAH therapies will be obsolete if the RV cannot recover. This suggests a need for studies to understand the mechanisms promoting dRV and cMFB regulation in the RV, which may be different than the left ventricle since the two ventricles have different embryologic origin (different heart fields), metabolism and biochemistry in the adult heart^14,15^.

Recent studies have identified a central role of mitochondrial calcium (mCa^++^) in the differentiation and activation of LV cMFB^16,17^. Uncoupling protein 2 (UCP2) has been shown to regulate the mitochondrial calcium uniporter (MCU) function and mitochondrial calcium (mCa^++^) uptake through mitochondrial calcium uptake 1 (MICU1)^18,19^. A reduction in UCP2—whether due to genetic knockout in mice or germline loss-of-function single nucleotide polymorphisms (SNPs) in humans—combined with increased MICU1 methylation, can result in decreased mCa^++^ levels. The resulting inhibition of Ca^++^-sensitive enzymes like Pyruvate Dehydrogenase (PDH) can alter the production of diffusible mitochondrial metabolites that can regulate epigenetic mechanisms in the nucleus, leading to a change of cell identity in cFB, toward cMFB. Our group has previously showed that genetic loss of UCP2 in mice promotes PAH and RV failure, and a UCP2 loss-of-function SNP (rs659366) is prevalent in PAH patients and associated with worse outcomes^20,21^. Furthermore, inflammatory cytokines like TNF-α increase from normal to cRV to dRV rats and are increased in the serum of PAH patients^22,23^. TNF-α inhibits UCP2 expression and decreases mCa^++24,25^, which could promote cMFB differentiation. In the present study, we show that a transition from cFB to cMFB contributes to RV decompensation in rodent and human PAH, driven by either genetic and/or or TNFα-induced UCP2 loss.

## Results

### Cardiac myofibroblasts (cMFB) and the transition from cRV to dRV in rats

We established a rat model of PAH using monocrotaline injection to study the hemodynamic transition from normal RV to cRV and ultimately to dRV. Although this inflammatory model does not replicate the lung histology of human PAH, it mirrors the rapid transition from cRV to dRV observed in patients. We studied rats 3 weeks post saline injection (CTRL) versus 3 weeks, and 5 weeks post monocrotaline injection. We used right heart catheterization (internal jugular canulation and advance of a Millar catheter tip into the main pulmonary artery), as well as echocardiography, to record mean PA pressure (i.e., the severity of PAH), RV systolic pressure (RVSP, i.e., the contractility of the right ventricle), right atrial (RA) pressure and cardiac output (i.e., the stage of heart failure), RV free wall thickness, RV end-diastolic diameter (RVEDD), and tricuspid annular plane systolic excursion (TAPSE) (i.e., the structural and functional RV remodeling)^26^. We defined cRV as a state of increased mean PA pressure (mPAP) and RVSP, but preserved cardiac output and mildly increased RA pressure; along with increased RV free wall thickness but not increased RVEDD. In contrast, dRV had high mPAP but a drop in RVSP, rate of contraction (dp/dt max), rate of relaxation (i.e., dp/dt min), cardiac output (CO) and TAPSE, as well as an increase in RA pressure and RVEDD (see methods for criterial) (**Figure 1A-C**). All the RVs and tissues labeled cRV or dRV thereafter, came from rats that strictly fulfilled these hemodynamic criteria. This offers an accurate distinction between the three RV functional stages.

**Figure 1.**
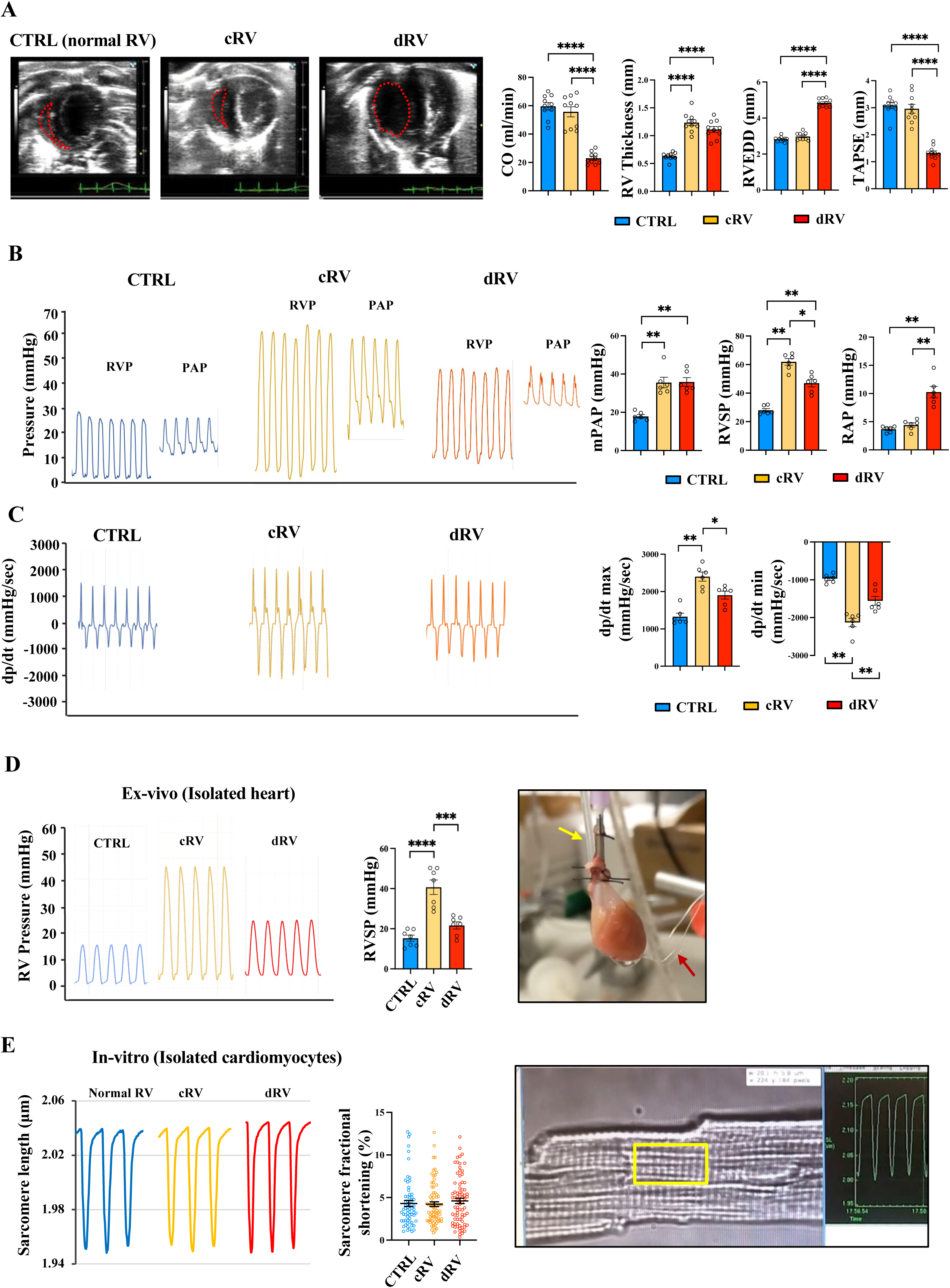
Right ventricular performance is reduced in decompensated RV in vivo and ex vivo, while cardiomyocyte contractility remains unchanged across control, cRV, and dRV. **A.** Representative echocardiography pictures (short axis) (left) show the size of RV chamber (red dotted line) in the three stages (Normal RV, cRV amd dRV) from PHT rats. Parameters measured by echocardiography (cardiac output, RV free wall thickness, RVEDD, TAPSE) are shown on the right, expressed as mean ± SEM; n = 10 animals for each group. ****P<0.0001. **B.** Representative tracings of RV (RVP) and pulmonary artery pressures (PAP) in CTRL, cRV and dRV from PHT rats (left) performed by close chest right heart catheterization. Parameters measured (mPAP, RVSP, RAP) are shown on the right and expressed as mean ± SEM; n = 6 animals for each group. *P<0.05, **P<0.01. **C.** Representative tracings of dp/dt of RV pressures in CTRL, cRV and dRV from PHT rats (left) with values (right) expressed as mean ± SEM; n = 6 animals for each group. *P<0.05, **P<0.01. **D.** Representative tracings (left) of RV pressures in ex-vivo isolated hearts from CTRL, cRV and dRV rats, with values (middle) expressed as mean ± SEM; n = 7 animals for each group. ***P<0.001. ****P<0.001. The picture (right) shows one isolated rat heart set up. A balloon was placed into RV (pointed by the yellow arrow) and pacemaker wires (pointed by the red arrow) were inserted into the myocardium. **E.** Representative tracings of sarcomere length (left) of RV cardiomyocyte contractility (sarcomere shortening) from normal RV, cRV and dRV from PHT rats with values (middle) expressed as mean ± SEM; n = ∼50 cells from 7 animals for each group. ***P<0.001. ****P<0.001. The picture (right) shows where the sarcomere shortening was measured (yellow rectangle). Comparisons were made using one-way ANOVA followed by Bonferroni post hoc analysis. **CTRL**, control (normal right ventricle, without monocrotaline injection); **cRV**, compensated right ventricle; **dRV**, decompensated right ventricle; **RVP**, right ventricle pressure; **PAP**, pulmonary artery pressure; **mPAP**, mean PA pressure; **RVSP**: right ventricular systolic pressure; **RAP**, right atrium pressure; **RVEDD**, right ventricular end-diastolic diameter; **TAPSE**, tricuspid annular plane systolic excursion.

Evaluating RV contractility in vivo is complicated by many factors, for example, afterload (e.g., mPAP), anesthesia or hormone effects. Therefore, we measured the RV contractility ex-vivo using the Langendorff perfusion system in which a balloon was inserted into the RV to measure contractility. Although we measured contractility without a pacemaker for the first 2 minutes, we also used a pacemaker beyond that point, as arrhythmias commonly occur, compromising our recordings and also to standardize measurements under the same heart rate (which tends to vary among different heart preparations). As expected, RV contractility decreased from cRV to dRV when studied under the same afterload **(Fig. 1D)**. However, when we isolated RV cardiomyocytes (CM) to study contractility in vitro (measured by sarcomere shortening), we found no significant differences among the three groups **(Fig. 1E)**. This suggests that the differences of contractility ex-vivo may be due to factors external to cardiomyocytes.

To investigate a potential role of cMFB, we first measured the number of RV cMFB which express both vimentin and α-SMA. We found that the number of cMFB is not different between normal RV and cRV, but it is dramatically increased in dRV **(Fig. 2A)**. We then isolated and cultured (myo)fibroblasts in short culture from the three groups and found that α-SMA is not present in normal RV and cRV fibroblasts (cFB) in contrast to dRV (**Supplementary Fig.1**). Furthermore, we also found that cMFB in dRV, in addition to an obvious change in cell shape compared to cFB, express high ki67 levels in vivo and in vitro compared with cFB, in keeping with their high proliferative nature **(Fig. 2B)**.

**Figure 2.**
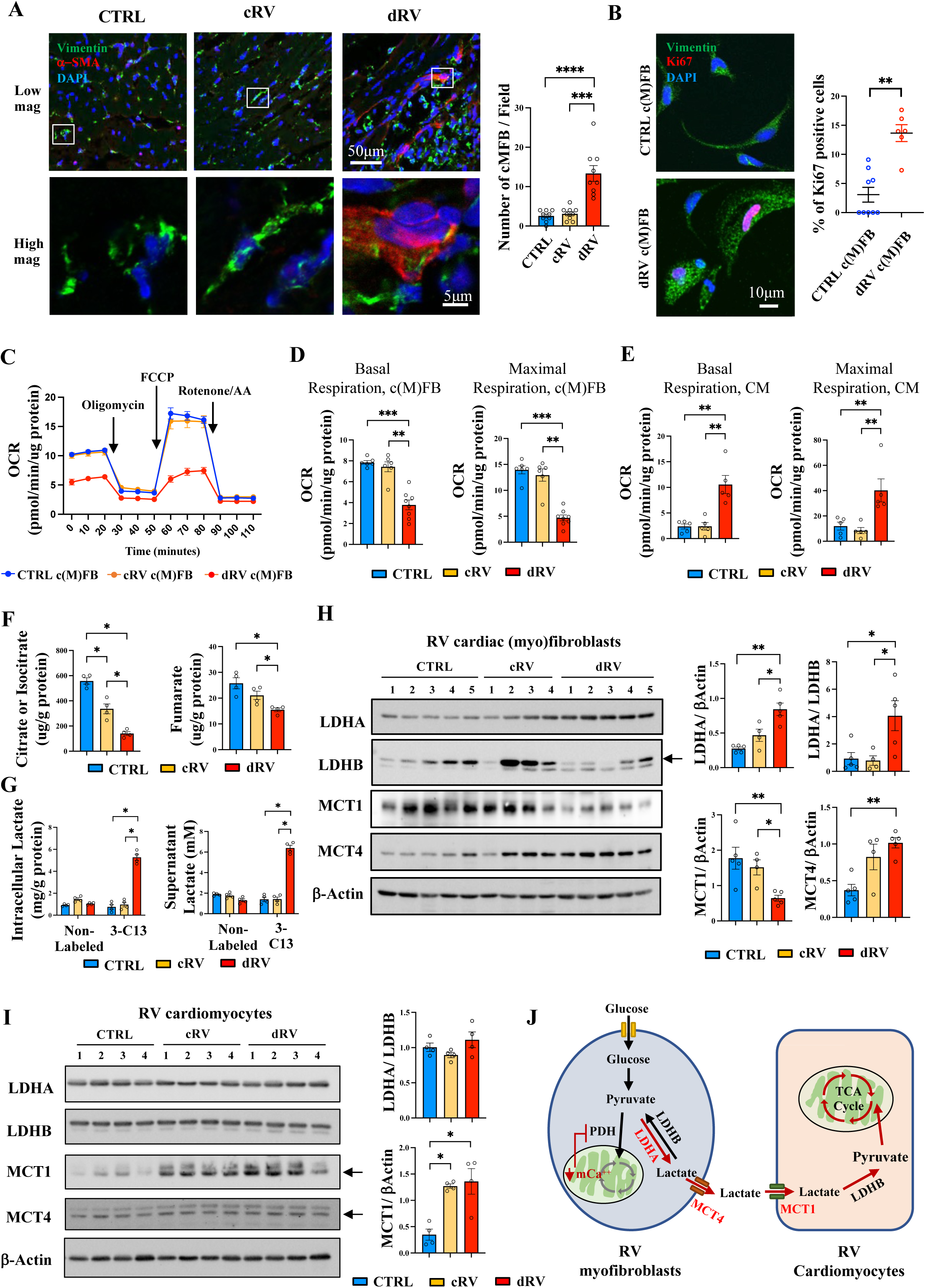
Decompensated RVs have more myofibroblasts, exhibiting metabolic cross talk with cardiomyocytes via lactate. **F.** Confocal microscopy (left) shows the number of myofibroblasts (positive with both α-SMA and Vimentin) in right ventricle frozen sections from Control, cRV and dRV from PHT rats, with values (right) expressed as mean ± SEM; n = 9 fields (∼ 200 fibroblasts from 3 animals for each group). ***P < 0.001; ****P < 0.0001. **G.** Confocal microscopy (left) and the ratio of Ki67 positive (with nuclear Ki67) cells in (myo)fibroblasts from normal RV and dRV from PHT rats (right) with values expressed as mean ± SEM; n = 9 fields (∼ 200 fibroblasts from 3 animals for each group). **P < 0.01. The comparison was made using the Mann-Whitney U test. **H.** A representative Seahorse experiment in (myo)fibroblasts from normal RV, cRV and dRV. Oligomycin inhibits ATP synthase and thus oxidative phosphorylation, while the ionophore FCCP uncouples the electron transport chain, exposing maximal respiration. The ETC complex I and III inhibitors rotenone and antimycin are used to uncover the mitochondrial component of respiration. So, the basal mitochondrial oxygen consumption rate (OCR) is the rate prior to any drugs, minus the residual rate after rotenone/antimycin. **I.** Seahorse values of mitochondrial basal and maximal OCR in cardiac (myo)fibroblasts from normal RV, cRV and dRV expressed as mean ± SEM; n = 6-8 from 3 animals for each group. **P <0.01; ***P < 0.001. **J.** Seahorse values mitochondrial basal and maximal OCR in cardiomyocytes from normal RV, cRV and dRV expressed as mean ± SEM; n =5 from 3 animals for each group. **P < 0.01. **K.** Mass Spectrometry shows the intracellular citrate levels in (myo)fibroblasts from normal RV, cRV and dRV with values expressed as mean ± SEM; n = 4 animals for each group. **P <0.01; ****P < 0.0001. **L.** Mass Spectrometry shows the intracellular and supernatant lactate levels in (myo)fibroblasts from normal RV, cRV after treatment with C-13 glucose (17.5mM) for 12h as mean ± SEM; n = 4 animals for each group. **P < 0.01; ****P < 0.0001. *3-C13: three Carbon-13 labeled lactate*. **M.** Immunoblot (left) shows the amount of LDHA, LDHB, MCT1, MCT4 and β-Actin in (myo)fibroblasts from normal RV, cRV and dRV. Quantification values (right) are expressed as mean ± SEM; n = 4-5 animals for each group *P < 0.05; **P < 0.01. **N.** Immunoblot shows (left) the levels of LDHA, LDHB, MCT1, MCT4 and β-Actin in cardiomyocytes from normal RV, cRV and dRV. Quantification values (right) are expressed as mean ± SEM; n = 4-5 animals for each group *P < 0.05; **P < 0.01. **O.** Cell-type-specific differences in the levels of pyruvate/lactate metabolizing enzymes and lactate importers/exporters, allow dRV cardiomyocytes to uptake the lactate produced by cMFB (as a result of their decreased mitochondrial respiration) and feed their TCA cycle to produce ATP, perhaps explaining their increased respiration compared to normal cardiomyocytes. Comparisons were made using Kruskal–Wallis tests followed by pairwise Mann–Whitney U tests. **CTRL**, control (normal right ventricle, without monocrotaline injection); **cRV**, compensated right ventricle; **dRV**, decompensated right ventricle; **α-SMA**, alpha smooth muscle actin; **DAPI**, 4′,6-diamidino-2-phenylindole (labels the nucleus); **Low mag**, low magnification; **High mag**, high magnification; **OCR**, oxygen consumption rate; **c(M)FB**, cardiac (myo)fibroblasts; **CM**, cardiomyocytes ; **LDH**, lactate dehydrogenase; **MCT**, monocarboxylate transporter.

We then studied whether there is a difference in mitochondrial function among the 3 different RV conditions using the Seahorse analyzer. We found that the dRV cMFB had decreased basal and maximal oxygen consumption rates (OCR) compared to normal RV and cRV cFB **(Fig. 2C-D)**. But an opposite pattern was found in CM, where the dRV CM had increased basal and maximal OCR **(Fig. 2E)**. Compared to normal and cRV cFB, dRV cMFB also had decreased TCA cycle metabolites, including citrate and fumarate, measured by mass spectrometry (**Fig. 2F**), which is in keeping with decreased glucose oxidation (GO). We then measured lactate, the terminal product of glycolysis, by using C-13 labeled glucose and found that dRV cMFB have increased levels (both intracellularly and extracellularly) of C-13-labeled lactate compared with normal and cRV cFB (**Fig. 2G**).

Intrigued by the disassociation between the mitochondrial function of the c(M)FB versus the CM during the progression of RV failure, we speculated that the two cells may communicate in a paracrine manner, in an attempt to harmonize energy homeostasis, similarly to what has been described between neurons and glial cells^27,28^. It is possible that the cardiomyocytes can use the glycolysis by-product lactate as a fuel to further support ATP synthesis. We speculated that the lactate produced by cMFB could be uptaken by CM, something that could explain the “paradoxical” increase in OCR that we found in CM. Therefore, we measured the levels of two enzymes: LDHA which preferentially converts pyruvate to lactate and LDHB which preferentially converts lactate to pyruvate; as well as two transporters of lactate, MCT4 for export and MCT1 for import of lactate. We found the LDHA progressively increased from normal RV to cRV to dRV c(M)FB, and the ratio of LDHA/LDHB sharply increased in dRV cMFB. Moreover, dRV cMFB had decreased MCT1 but increased MCT4 compared with normal RV and cRV cFB (**Fig. 2H**). In contrast, dRV CM had increased MCT1 compared with normal CM (**Fig. 2I**). Taken together, these data suggest that dRV cMFB could produce and secrete lactate which could be taken up by dRV CM. To prove that lactate can be used as a fuel to generate ATP in CM, we treated CM from normal RV and dRV with C-13 labeled lactate and then measured C-13 labeled TCA cycle intermediates. We found that the dRV CM had increased C-13-labeled metabolites compared to normal CM (**Supplementary Fig. 2**). The proposed mechanism of the metabolic crosstalk between cMFB and CM is shown in **Fig. 2J**.

The OCR is a reflection of the overall mitochondrial function, particularly the production of NADH and FADH from the TCA cycle, which are used in Electron Transport Chain to generate ATP along with oxygen. The production of NADH and FADH depends on the activity of many enzymes that are mCa^++^-dependent (e.g., PDH). Thus, we measured mCa^++^ in c(M)FB and found decreased mCa^++^ from normal to cRV and a further decrease to dRV cMFB **(Fig. 3A)**. We then measured PDH activity and as shown in **Fig. 3B**, the dRV cMFB had decreased PDH activity compared to those from normal and cRV.

**Figure 3.**
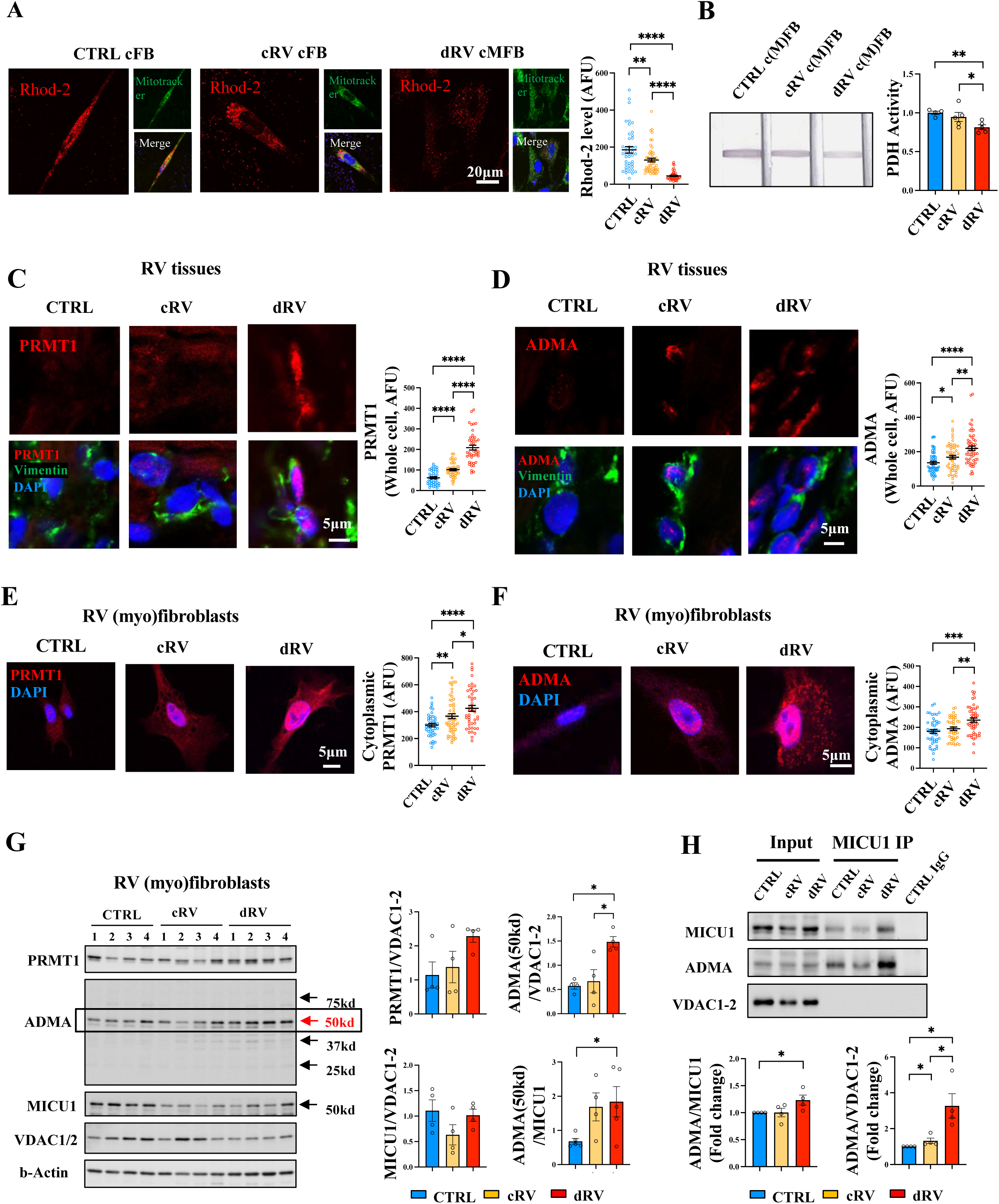
Mitochondrial calcium levels are decreased and MICU1 methylation is increased in dRV myofibroblasts compared with normal RV and cRV fibroblasts. **A.** Confocal microscopy (left) showing mitochondrial calcium level (Rhod-2 fluorescence) in cultured (myo)fibroblasts from normal RV, cRV and dRV. Quantification values (right) are expressed as mean ± SEM; n = 50 cells from 3 animals for each group. **P < 0.01; ****P < 0.0001. **B.** PDH activity, measured by an NADH-based enzymatic activity ElLISA, in (myo)fibroblasts from normal RV (CTRL), cRV and dRV. Quantification values (right) are expressed as mean ± SEM; n = 50 cells from 3 experiments. n = 4 animals for each group. *P < 0.05; **P < 0.01. **C.** Confocal microscope (left) shows the PRMT1 (i.e. arginine methyltransferase) level in (myo)fibroblasts (vimentin-positive cells) from right ventricle frozen sections of Control, cRV and dRV rats. Quantification values (right) are expressed as mean ± SEM; n = 50 cells from 3 animals for each group. ****P < 0.0001. **D.** Confocal microscopy (left) shows the AMDA level (i.e. arginine methylation level) in (myo)fibroblasts (vimentin-positive cells) from Control, cRV and dRV tissues. Quantification values (right) are expressed as mean ± SEM; n = 50 cells from 3 animals for each group. *P < 0.05; **P < 0.01; ****P < 0.0001. **E.** Confocal microscopy (left) shows the cytoplasmic PRMT1 level in cultured cardiac (myo)fibroblasts from normal RV, cRV and dRVs. Quantification values (right) are expressed as mean ± SEM; n = 50 cells from 3 experiments. *P < 0.05; **P < 0.01; ****P < 0.0001. **F.** Confocal microscopy (left) shows the cytoplasmic ADMA levels in cultured (myo)fibroblasts from normal RV, cRV and dRVs. Quantification values (right) are expressed as mean ± SEM; n = 50 cells from 3 experiments. ***P < 0.001. **G.** Immunoblot (left) of PRMT1, ADMA, MICU1, VDAC1/2 and β-Actin in (myo)fibroblasts from normal RV, cRV and dRV of rats. Quantification values (right) are expressed as mean ± SEM; n = 4 animals for each group. *P < 0.05. **H.** MICU1 Immunoprecipitation (above) on isolated mitochondria from (myo)fibroblasts shows the amount of MICU1 and ADMA. MICU1 methylation is shown as ADMA/MICU1. Quantification values (below) are expressed as mean ± SEM; n = 4 animals for each group. **A, C, D, E, F comparisons** were made using one-way ANOVA followed by Bonferroni post hoc analysis; **B, G, H** were made using Kruskal–Wallis tests followed by pairwise Mann–Whitney U tests. **CTRL**, control (normal right ventricle, without monocrotaline injection); **cRV**, compensated right ventricle; **dRV**, decompensated right ventricle; **cFB,** cardiac fibroblasts; **cMFB,** cardiac myofibroblasts; **c(M)FB**, cardiac (myo)fibroblasts; **PDH**, pyruvate dehydrogenase; **PRMT1**, Protein Arginine Methyltransferase 1; **ADMA**, Asymmetric dimethylarginine; **DAPI**, 4′,6-diamidino-2-phenylindole (labels the nucleus); **MICU1**, Mitochondrial Calcium Uptake 1; **VDAC**, Voltage-Dependent Anion Channels; **IP**, Immunoprecipitation.

To determine the basis of this difference, we first measure the levels of the mCa^++^ uniporter MCU and its two main components, MICU1 and UCP2. In addition, because the function of the MICU1 has been shown to depend on its methylation by the methyltransferase PRMT1, we also measured the expression of PRMT and MICU methylation status using ADMA and MICU immunoprecipitation. First, with RV tissue staining, we found that the cytoplasmic PRMT1 and ADMA levels are increased from normal to cRV cFB and are further increased in dRV cMFB (**Fig. 3C-F**).). Similar results were found in immunoblots, in which the ADMA is increased in dRV cMFB compared with normal RV and cRV cFB **(Fig. 3G).** The PRMT1 had a trend to increase in dRV cMFB compared with normal and cRV cFB, but there was no significant difference. To measure the methylation status of MICU1 specifically, we isolated mitochondria from c(M)FB, performed a MICU1 immunoprecipitation, and measured the level of ADMA. We found MICU1 methylation is increased in dRV cMFB compared to normal RV and cRV cFB **(Fig. 3H)**. There was no difference in MCU protein level in c(M)FB among the three groups, suggesting that the decreased mCa^++^ is not due to MCU levels change, but, rather, an increase in the MICU1 methylation **(Supplementary Fig. 3)**. In contrast, we found no difference in PRMT1 and ADMA in CM among the three RV groups **(Supplementary Fig. 4),** which means this change is fibroblasts-specific.

UCP2 level were exclusively decreased in dRV tissue compared with cRV and normal RV, while this decrease was not found in the corresponding LVs, suggesting that is RV-specific **(Fig. 4A)**. The UCP2 level was decreased in dRV c(M)FB compared to normal RV and cRV (**Fig. 4B**), but not in CM (**Supplementary Fig. 5A-B**), meaning that fibroblasts are the main source of the reduced UCP2 levels observed in the RV.

**Figure 4.**
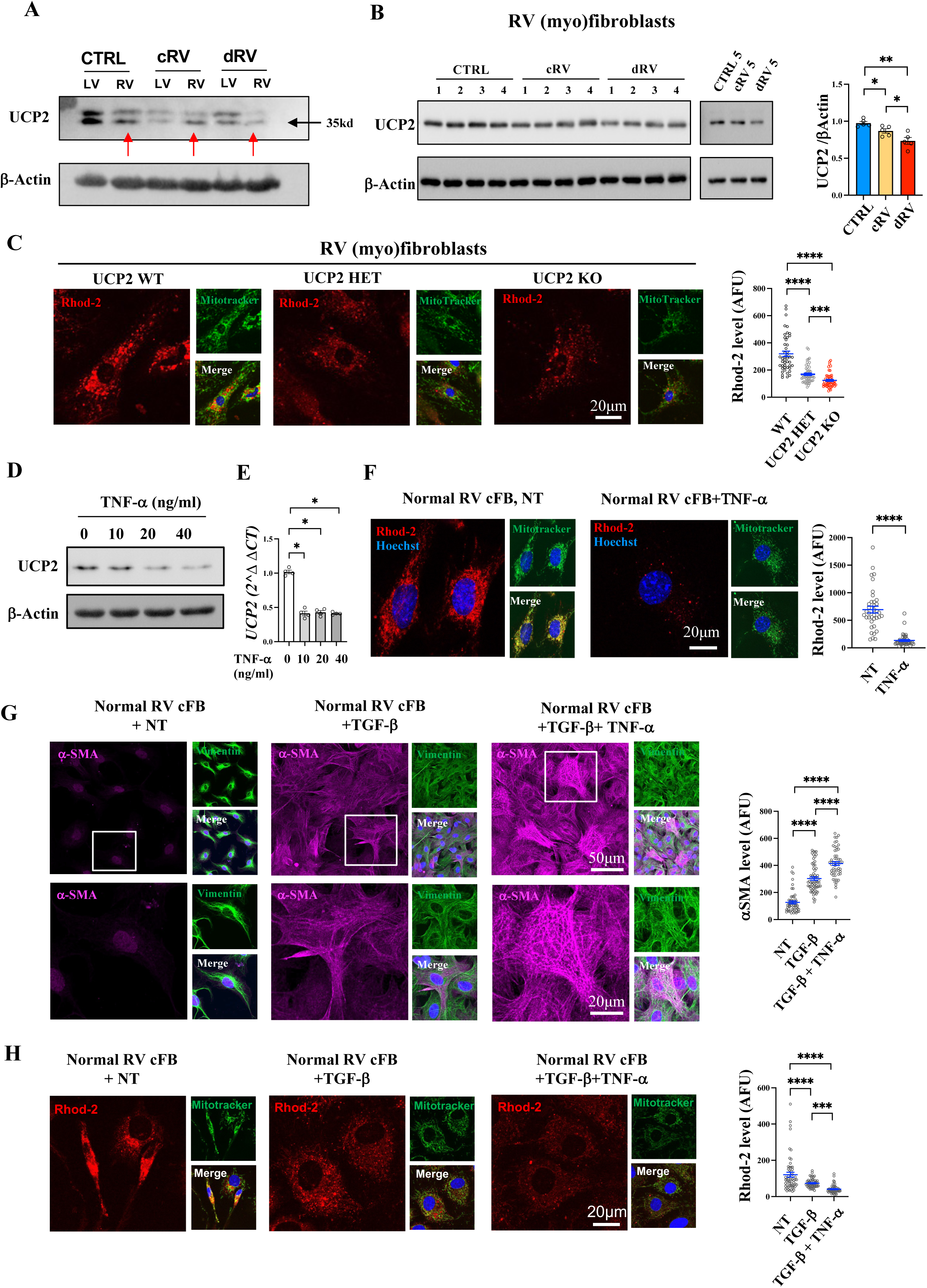
UCP2 levels decrease from control to cRV to dRV (myo)fibroblasts in vivo, while TNF-*α* reduces UCP2 expression and mitochondrial calcium levels, promoting fibroblast activation in vitro. **P.** Immunoblot shows the UCP2 protein level in the myocardium of the left ventricle and right ventricles from Control, cRV and dRV PHT rats (red arrows point the right ventricles). UCP2 is the lower band with a molecular weight of 35kd (verified by UCP2 WT and KO immunoblot). **Q.** Immunoblot shows the UCP2 protein level in right ventricular (myo)fibroblasts from normal RV, cRV and dRV. Quantification values (**right**) normalized by b-Actin are expressed as mean ± SEM; n = 5 animals for each group. *P < 0.05; **P < 0.01. **R.** Confocal microscopy (left) shows the mitochondrial calcium level (Rhod-2 fluorescence) in cultured right ventricular (myo)fibroblasts from WT, UCP2 heterozygous and UCP2 knockout mice. Quantification values (right) are expressed as mean ± SEM; n = 50 cells from 3 experiments. ***P < 0.001; ****P < 0.0001. **S.** Immunoblot shows the UCP2 protein levels in normal right ventricular fibroblasts treated with or without TNF-α at doses of 10, 20, or 40 ng/ml. **T.** q-PCR shows the UCP2 RNA levels in normal right ventricular fibroblasts treated with or without TNF-α at doses of 10, 20, or 40 ng/ml. **U.** Confocal microscopy (left) shows the mitochondrial calcium level (Rhod-2 fluorescence) in cultured normal right ventricular fibroblasts treated with or without TNF-α at a doses of 20 ng/ml. Quantification values (right) are expressed as mean ± SEM; n = 50 cells from 3 experiments. ****P < 0.0001. The comparison was made using unpaired Student t test. **V.** Confocal microscopy (left) shows the α-SMA level (far red) in cultured normal right ventricular fibroblasts treated with vehicle (NT), TGF-β or TGFβ+TNF-α. Quantification values (right) are expressed as mean ± SEM; n = 50 cells from 3 experiments. ****P < 0.0001. **W.** Confocal microscopy (left) shows the mitochondrial calcium level (Rhod-2 fluorescence) in cultured normal right ventricular fibroblasts treated with vehicle (NT), TGF-β or TGF-β+TNF-α. Quantification values (right) are expressed as mean ± SEM; n = 50 cells from 3 experiments. ***P < 0.001, ****P < 0.0001. **C, F, G, H** were made using one-way ANOVA followed by Bonferroni post hoc analysis; **B, E** were made using Kruskal–Wallis tests followed by pairwise Mann–Whitney U tests. **CTRL**, control (normal right ventricle, without monocrotaline injection); **cRV**, compensated right ventricle; **dRV**, decompensated right ventricle; **LV**, left ventricle; **RV**, right ventricle; **UCP2**, uncoupling Protein 2; **WT**, wild type; **HET**, heterozygous; **KO**, knockout; **cFB**, cardiac fibroblasts; **NT**, no treatment; **α-SMA**, alpha smooth muscle actin; **TGF-β**, Transforming growth factor-beta; **TNF-α**, tumor necrosis factor alpha.

To determine the role of UCP2 in the regulation of mCa^++^ independent of M1CU1 methylation, we measured mCa^++^ level in cFB from wild-type, UCP2 heterozygote and homozygote KO mice (KO was confirmed in both RNA and protein levels in cMFB in **Supplementary Fig. 6A-B)**. We found a gene dose-dependent decrease in mCa^++^ as UCP2 levels decreased **(Fig. 4C)**. These data together suggest that the reason for the progressive loss of mCa^++^ from normal to cRV to dRV may be an increase of MICU1 methylation combined with a loss of UCP2 in cMFB. This can explain the increased number of cMFB from cRV to dRV.

Patients with PAH have a very inflammatory phenotype compared other types of pulmonary hypertension, and we have previously shown that in the transition from cRV to dRV in rats, there is an increase in the cytokine TNF-α^22^. Since TNF-α can decrease the expression of UCP2^29^, we measure the UCP2 level in fibroblasts under different doses of TNF-α. We found that UCP2 mRNA and protein levels decreased in a dose-dependent manner **(Fig. 4D, E)**. Not surprisingly, the mCa^++^ level also decreased after TNF-α treatment **(Fig. 4F)**. To determine whether TNF-α increases α-SMA over and above the classic inducer of cMFB (i.e., TGF-β), we studied the effect of TNF-α on top of TGF-β in mCa^++^ and αSNA levels in RV fibroblasts. Compared to vehicle, cFB treated with TGF-β alone have increased α-SMA and decreased mCa^++^; and cFB treated with TGF-β and TNF-α show further increased α-SMA and decreased mCa^++^ **(Fig. 4G-H)**. Because the TGF-β axis is disturbed in PAH due to germline mutations or other factors^30–35^, our data suggest that in addition to the pulmonary vascular cells, the RV fibroblasts may also be affected, contributing to dRV, particularly in patients with scleroderma.

### RV cMFB and UCP2 in patients with PAH and secondary PHT

We studied two cohorts of patients with PAH (cohort 1 from Laval University and cohort 2 from the University of Alberta) and one cohort of patients with secondary pulmonary hypertension (sPHT, from Duke University). Cohort 1 included RV tissues (from heart/lung transplant recipients or autopsies) from 30 patients without (sudden cardiac death with no known heart disease and macroscopically normal RVs) or with group 1 PAH (**supplementary table 1**), where the PAH diagnosis had been confirmed with right heart catheterization (RHC) and echocardiography. The vast majority of PAH patients had available mPAP, cardiac index (CI) and TAPSE values. Cohort 2 included 21 patients with PAH (**supplementary table 2**) who underwent RHC and echocardiography within 2 days from each other, approximately one year after the establishment of diagnosis and initiation of double oral PAH therapy with phosphodiesterase type 5 inhibitors and endothelin receptor antagonists. Cohort 2 did not have RV tissues, as they were all alive and not transplanted, but offered blood. The same treatment protocol allowed us to compare cRV and dRV independent of therapy. The diagnosis of PAH in both cohorts was based on RHC mPAP >20mmHg and PAWP<13 mmHg, along with structurally and functionally normal LV on ECHO and the absence of thromboembolic and parenchymal lung disease (chest CT, pulmonary function tests). Cohort 3 included 30 patients whose RVs were obtained from heart transplant recipients for ischemic or non-ischemic cardiomyopathy (**supplementary table 3**). 21 of them had sPHT (mPAP>20mmHg) with available TAPSE and/or CI values for most patients.

We identified dRV as CI<2.2 L/min/m^2^ and/or TAPSE<1.5 cm for all cohorts. We studied RVs using immunohistochemistry, droplet digital PCR (ddPCR), immunoblots; and the presence of a heterozygous and homozygous loss-of-function UCP2 SNP (rs659366) by ddPCR in either RV tissues or blood cells. We correlated the number of RV cMFB or the presence of the SNP with RV UCP2 mRNA and protein levels as well as with CI and TAPSE.

**1.** PAH cohort (cohort 1 and 2): We found that the number of cMFB did not increase from normal RV to cRV but increased sharply in dRV in cohort 1 (**Fig. 5A**); and correlated negatively with TAPSE and CI (**Fig. 5B-C**). We also found the cRV cFB have increased cytoplasmic PRMT1 and ADMA levels, compared to normal RV fibroblasts but not significantly different compared to dRV cMFB (**Fig. 5D-E**); and no correlation was found with TAPSE or CI (Supplementary Fig. 7). In contrast, the cMFB UCP2 levels are decreased in dRV compared to normal and cRV fibroblasts (**Fig. 6A**). The UCP2 protein and mRNA levels correlated negatively with the number of RV cMFB and positively with TAPSE or CI (**Fig. 6B-G**). These human data suggested that that the main difference from cRV to dRV is the progressive decrease in UCP2 levels, similarly to the rat data.

**Figure 5.**
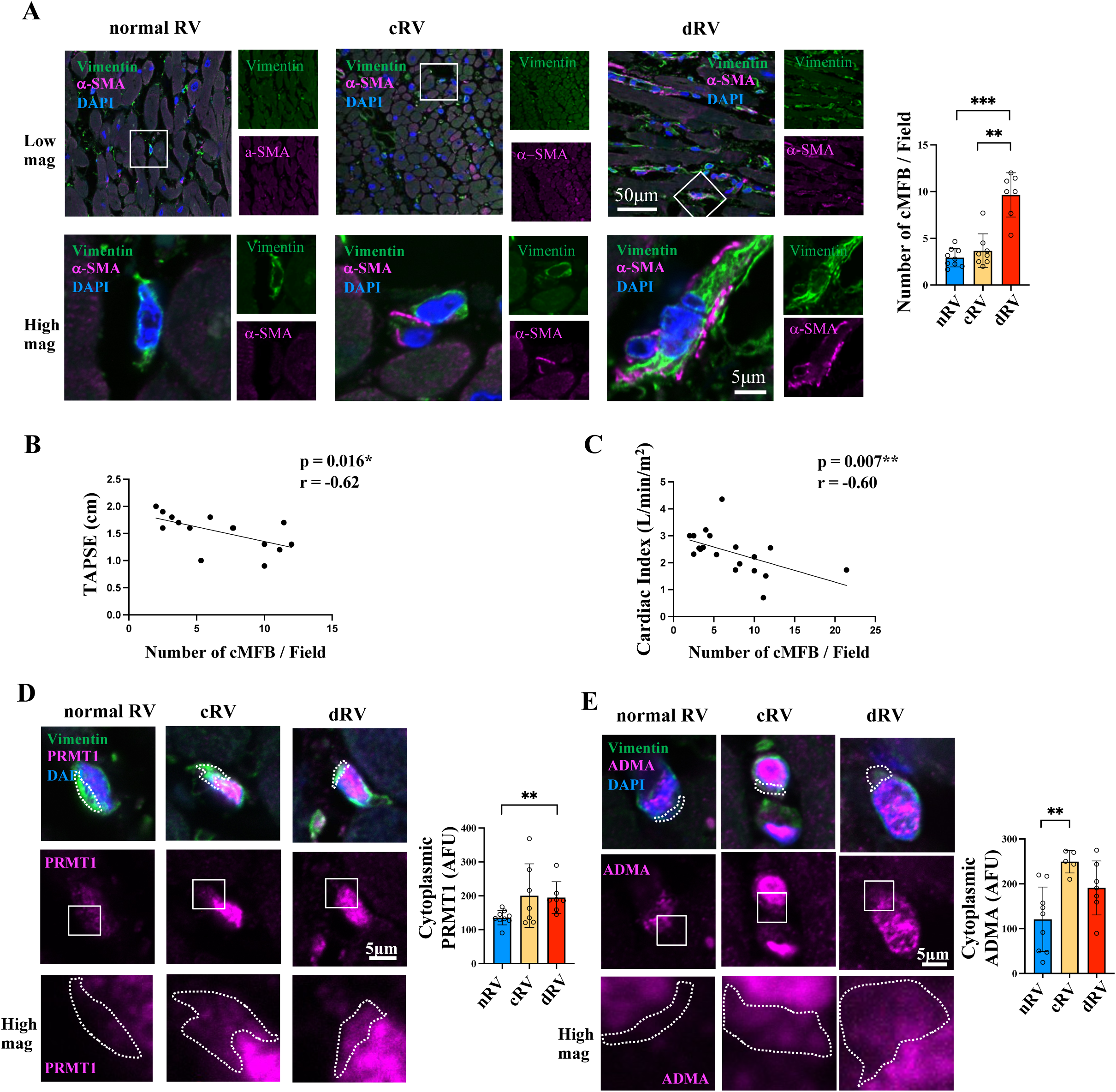
Myofibroblasts are activated in human decompensated RV, while arginine methylation in (myo)fibroblasts increases in compensated RV but does not further increase in decompensated RV compared with control (normal RV). **A.** Confocal microscopy (left) shows the number of myofibroblasts (positive with both α-SMA and Vimentin) from paraffin-embedded right ventricle sections from CTRL (n=9), cRV (n=10) and dRV (n=11) cohort 1patients with PAH. Scatter dot plots represent individual values. Quantification values (right) are expressed as mean ± SEM; ****P < 0.0001. **B.** Spearman correlation test shows the correlation between number of myofibroblasts with TAPSE in patients with cohort 1 PAH. Scatter dot plots represent individual values. *p<0.05 **C.** Spearman correlation test shows the correlation between number of myofibroblasts with Cardiac Index in patients with cohort 1 PAH. Scatter dot plots represent individual values. *p<0.05 **D.** Confocal microscopy (left) shows the cytoplasmic PRMT1 level in (myo)fibroblasts (vimentin-positive cells) from paraffin-embedded right ventricle sections from CTRL (n=9), cRV (n=10) and dRV (n=11) cohort 1 patients with PAH. The dotted line tracing follows the Vimentin signal in order to trace the cytoplasm. Scatter dot plots represent individual values. Quantification values (right) are expressed as mean ± SEM; *P < 0.05; ****P < 0.0001. **E.** Confocal microscope (left) shows the cytoplasmic ADMA level in (myo)fibroblasts (vimentin-positive cells) from paraffin-embedded right ventricle sections from CTRL (n=9), cRV (n=10) and dRV (n=11) cohort 1 patients with PAH. The dotted line tracing follows the Vimentin signal in order to trace the cytoplasm. Scatter dot plots represent individual values. Quantification values (right) are expressed as mean ± SEM; **P < 0.01; ***P < 0.001. **A, D, E** were made using Kruskal–Wallis tests followed by pairwise Mann–Whitney U tests. **nRV,** normal right ventricle; **cRV**, compensated right ventricle; **dRV**, decompensated right ventricle; **RV**, right ventricle; **α-SMA**, alpha smooth muscle actin; **DAPI**, 4′,6-diamidino-2-phenylindole (labels the nucleus); **PRMT1**, Protein Arginine Methyltransferase 1; **ADMA**, Asymmetric dimethylarginine.

**Figure 6.**
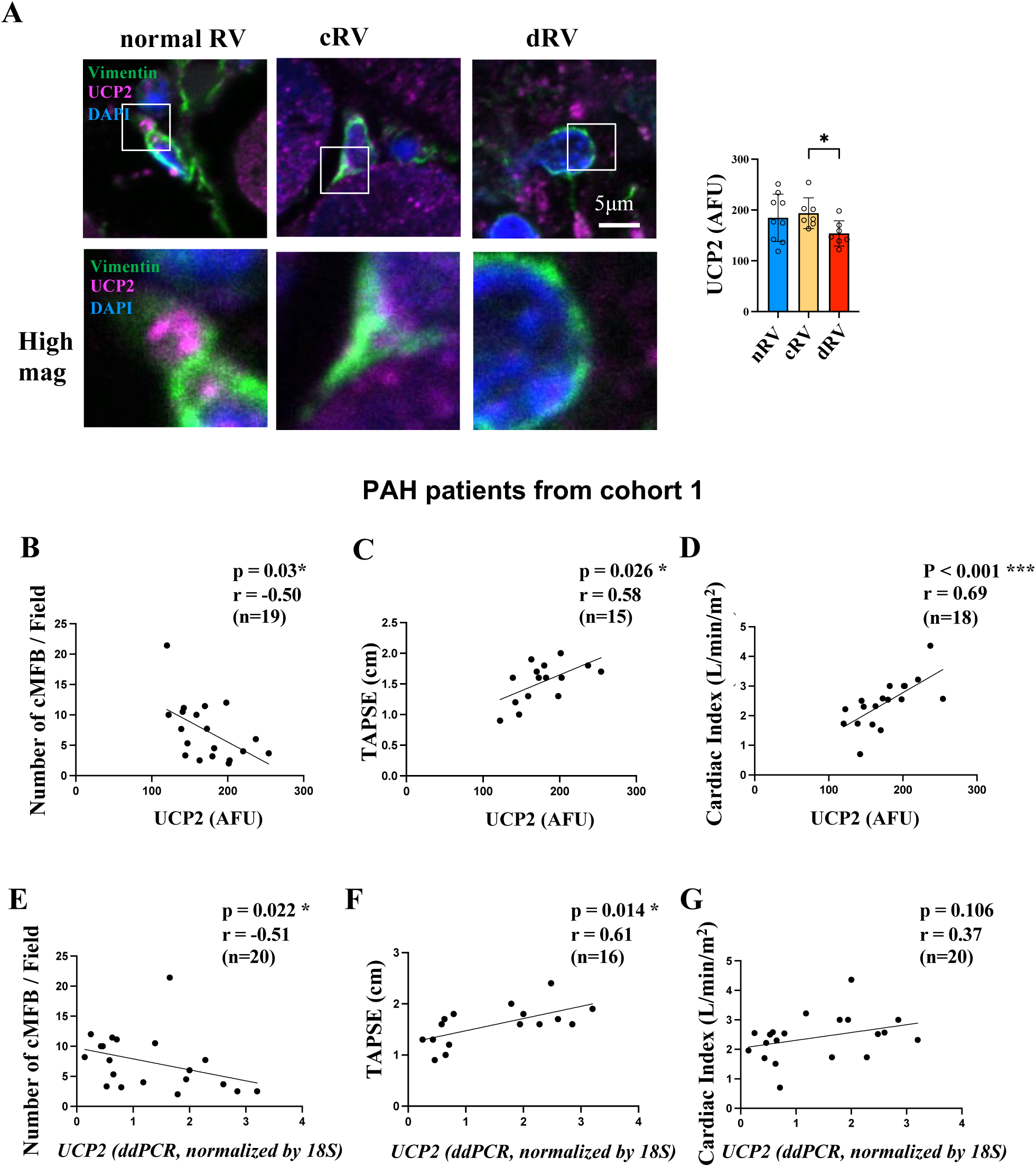
UCP2 levels correlate negatively with the number of cardiac myofibroblasts and positively with TAPSE and cardiac index in PAH patients. **A.** Confocal microscope (left) shows the UCP2 level in (myo)fibroblasts (vimentin-positive cells) from paraffin-embedded right ventricle sections from CTRL (n=9), cRV (n=10) and dRV (n=11) cohort 1 patients with PAH. Scatter dot plots represent individual values. Quantification values (right) are expressed as mean ± SEM; *P < 0.05; ****P < 0.0001. This comparison was made using Kruskal–Wallis tests followed by pairwise Mann– Whitney U tests. **B.** Spearman correlation test shows the correlation between UCP2 immunofluorescence intensity in (myo)fibroblasts with number of myofibroblasts/field in cohort 1 patients with PAH. Scatter dot plots represent individual values. *p<0.05 **C.** Spearman correlation test shows the correlation between UCP2 immunofluorescence intensity in (myo)fibroblasts with TAPSE cohort 1 patients with PAH. Scatter dot plots represent individual values. *p<0.05 **D.** Spearman correlation test shows the correlation between UCP2 immunofluorescence intensity in (myo)fibroblasts with Cardiac Index in cohort 1 patients with PAH. Scatter dot plots represent individual values. **p<0.01 **E.** Spearman correlation test shows the correlation between UCP2 RNA level with number of myofibroblasts/field in patients with PAH. Scatter dot plots represent individual values. *p<0.05 **F.** Spearman correlation test shows the correlation between UCP2 RNA level with TAPSE in cohort 1 patients with PAH. Scatter dot plots show individual values. *p<0.05 **G.** Spearman correlation test shows the correlation between UCP2 RNA level with Cardiac Index (CI) cohort 1 patients with PAH. Scatter dot plots represent individual values. **nRV,** normal right ventricle; **cRV,** compensated right ventricle; **dRV**, decompensated right ventricle; **UCP2**, uncoupling protein 2; **DAPI**, 4′,6-diamidino-2-phenylindole (labels the nucleus); **TAPSE**, tricuspid annular plane systolic excursion; **PAH**, pulmonary arterial hypertension; **UCP2,** uncoupling protein 2.

The UCP2 SNP (rs659366) affects the promoter of the UCP2 gene, resulting in decreased UCP2 mRNA levels in the -866 A allele (UCP2 SNP) compared to GG allele (WT)^28^. The presence of this loss-of-function UCP2 SNP is associated with worse outcomes in PAH patients^19^. We measured the presence of UCP2 SNP from all human RV tissues (cohorts 1 and 3) and found that the presence of the SNPs is associated with decreased UCP2 protein and mRNA levels in a gene dose-dependent manner, i.e., progressively decreasing from wild-type (WT) to heterozygotes (HET) to homozygotes (HOM) (**Fig. 7A-B**). In cohort 1 patients with PAH, TAPSE was associated with the presence of UCP2 SNP and there was a trend for CI. This was true even for patients that had similar mPAP, i.e., patients with WT UCP2 (white dots in Fig 7C) had higher TAPSE than patients with heterozygous SNP (grey dots) and homozygous UCP2 SNP (red dots), even though they had similar mPAPs (**Fig. 7 C-D**). In cohort 2 patients with PAH, the same associations were found but with significance in CI (**Fig. 7 E-F**). When we pulled cohort 1 and 2 patients with PAH, achieving sample sizes of 36 and 39 for TAPSE and CI respectively, we found that the presence of the SNP clearly separated dRV from cRV based on both TAPSE and CI (**Fig. 7 G-H**). In these patients, the proportion of patients with heterozygous or homozygous SNP increased from 39.1 % of cRV patients to 81.3 % of patients with dRV (**Fig. 7 I**).

**Figure 7.**
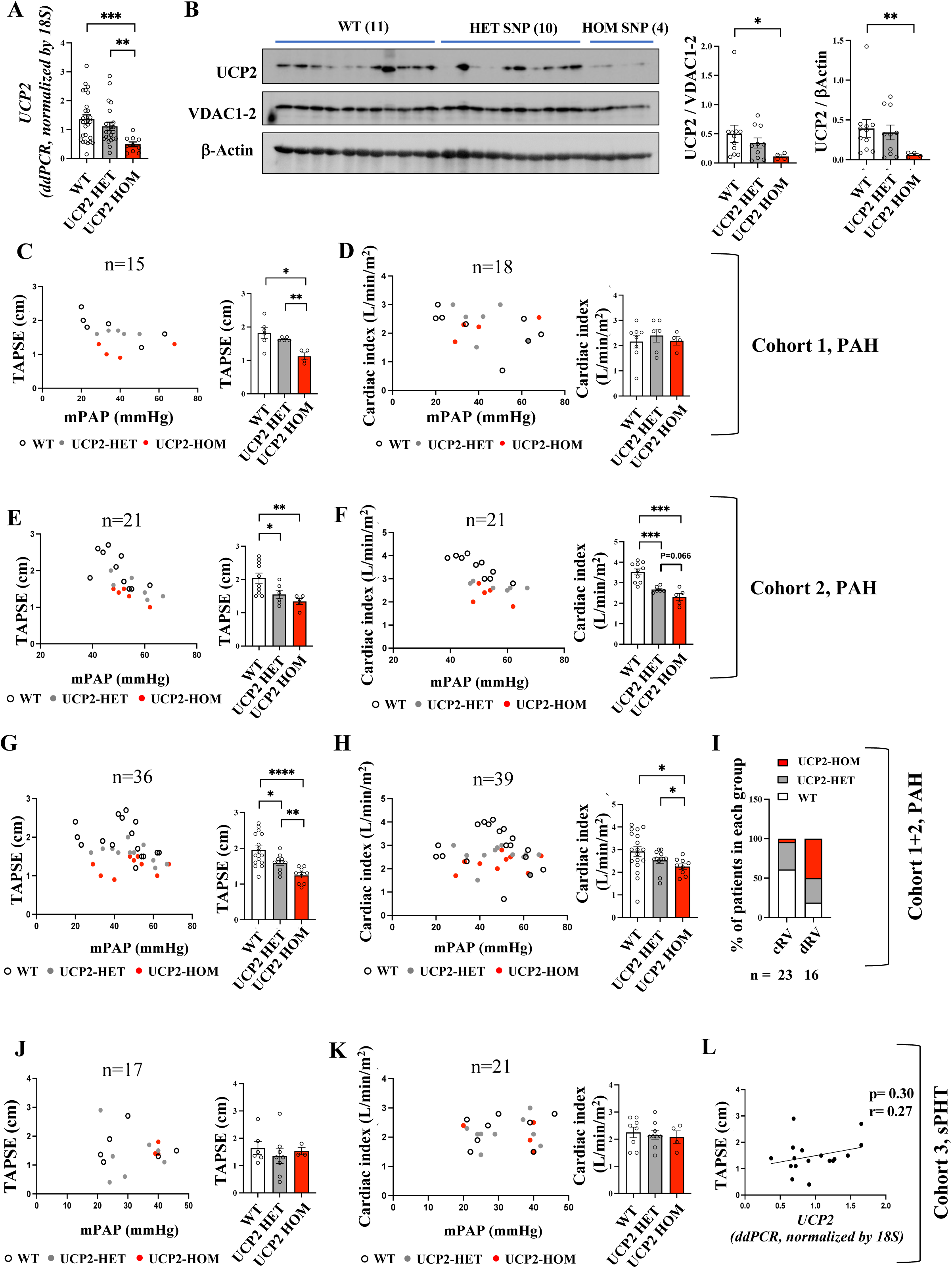
UCP2 SNP, reflecting UCP2 RNA and protein levels, correlates with TAPSE and cardiac index in patients with PAH but not in those with sPHT. **A.** qPCR shows the UCP2 RNA level in human RV (from cohort 1and 3) genotyped with WT, UCP2 HET and HOM SNP. Scatter dot plots represent individual values. Quantification values are expressed as mean ± SEM. ***P < 0.001. **B.** Western blot (left) shows the UCP2 protein level in human RV genotyped with WT, UCP2 HET and HOM SNP. Scatter dot plots represent individual values. Quantification values (right) are expressed as mean ± SEM; *P < 0.05. **C.** UCP2 genotyping and TAPSE in patients with PAH (mPAP>20mmHg from cohort 1). Patients were genotyped for UCP2 and categorized into WT (n=6), UCP2 HET SNP (n=5) and UCP2 HOM SNP (n=4). Scatter dot plots represent individual values. Quantification values (right) are expressed as mean ± SEM *P < 0.05; **P < 0.01. **D.** UCP2 genotyping and echocardiography show the UCP2 genotype and Cardiac Index in patients with PAH (mPAP>20mmHg from cohort 1), sub-grouped by UCP2 genotype: WT (n=8), UCP2 HET SNP (n=6) and UCP2 HOM SNP (n=4). Scatter dot plots represent individual values. Quantification values (right) are expressed as mean ± SEM. **E.** UCP2 genotyping and echocardiography show the UCP2 genotype (blood) and Cardiac index in patients with PAH (mPAP>20mmHg from cohort 2). Patients were genotyped for UCP2 and categorized into WT (n=10), UCP2 HET SNP (n=6) and UCP2 HOM SNP (n=5). Scatter dot plots represent individual values. Quantification values (**right**) are expressed as mean ± SEM. *P < 0.05; **P < 0.01. **F.** UCP2 genotyping and echocardiography show the UCP2 genotype and Cardiac Index in patients with PAH (mPAP>20mmHg from cohort 2). Patients were genotyped for UCP2 and categorized into WT (n=10), UCP2 HET SNP (n=6) and UCP2 HOM SNP (n=5). Scatter dot plots represent individual values. Quantification values (**right**) are expressed as mean ± SEM. **G.** UCP2 genotyping and echocardiography show the UCP2 genotype and TAPSE in patients with PAH (mPAP>20mmHg from cohort 1and 2). Patients were genotyped for UCP2 and categorized into WT (n=16), UCP2 HET SNP (n=11) and UCP2 HOM SNP (n=9). Scatter dot plots represent individual values. Quantification values (right) are expressed as mean ± SEM *P < 0.05; **P < 0.01; ***P < 0.001. **H.** UCP2 genotyping and echocardiography show the UCP2 genotype and Cardiac index in patients with PAH (mPAP>20mmHg from cohort 1and 2). Patients were genotyped for UCP2 and categorized into WT (n=18), UCP2 HET SNP (n=12) and UCP2 HOM SNP (n=9). Scatter dot plots represent individual values. Quantification values (right) are expressed as mean ± SEM. *P < 0.05. **I.** UCP2 genotyping shows the ratio of patients with WT, UCP2-HET SNP and UCP2-HOM SNP in cRV vs dRV in patients with PAH (Cohort 1+2). **J.** UCP2 genotype and TAPSE in patients with sPHT (mPAP>20 mmHg from Cohort 3). Patients were genotyped for UCP2 and categorized into WT (n=6), UCP2 HET SNP (n=8) and UCP2 HOM SNP (n=3). Scatter dot plots represent individual values. Quantification values (**right**) are expressed as mean ± SEM. **K.** UCP2 genotype and TAPSE in patients with with sPHT (mPAP>20 mmHg from Cohort 3). Patients were genotyped for UCP2 and categorized into WT (n=8), UCP2 HET SNP (n=9) and UCP2 HOM SNP (n=4). Scatter dot plots represent individual values. Quantification values (**right**) are expressed as mean ± SEM. **L.** Spearman correlation test shows lack of orrelation between UCP2 RNA level and TAPSE in patients with sPHT (mPAP>20 mmHg from Cohort 3). Scatter dot plots show individual values. These comparisons were made using Kruskal–Wallis tests followed by pairwise Mann–Whitney U tests. **SNP,** Single Nucleotide Polymorphism; **UCP2**, uncoupling protein 2; **VDAC**, Voltage-Dependent Anion Channels; **WT,** wild type; **UCP2 HET,** heterozygous UCP2 SNP; **UCP2 HOM,** homozygous UCP2 SNP; **TAPSE,** Tricuspid Annular Plane Systolic Excursion; **mPAP:** mean PA pressure; **PAH,** pulmonary arterial hypertension; s**PHT**: Secondary pulmonary hypertension; **nRV,** normal right ventricle; **cRV,** compensated right ventricle; **dRV**, decompensated right ventricle; **cFB,** cardiac fibroblasts; **cMFB,** cardiac myofibroblasts.

**2. sPHT (cohort 3).** In contrast to the PAH cohorts, no differences in the number of RV cardiac myofibroblasts (cMFBs) were observed among normal, cRV, and dRV in the sPHT cohort (**Supplementary Fig. 8A**). However, the sPHT RVs had higher numbers of cMFBs compared to control patients in cohort 1 studied under identical immunohistochemistry protocols. No significant differences were found in cytoplasmic PRMT1 or UCP2 levels in RV c(M)FBs in sPHT RVs (**Supplementary Fig. 8B, C**). Predictably, in cohort 3 patients with pulmonary hypertension (i.e., mPAP>20 mmHg), we observed no association between TAPSE or CI and the presence of the UCP2 SNP; or with the levels of RV UCP2 mRNA levels (**Fig. 7 J-L).** There are several potential explanations for this difference, in addition to the fact that the sPHT mPAPs were lower than in the PAH cohorts. First, sPHT is generally associated with a lower inflammatory burden compared to PAH and our rat data showed that TNF-α decreases the expression of UCP2. Second, in ischemic cardiomyopathy which accounted for many cases in cohort 3, RV dysfunction may primarily result from ischemia due to right coronary artery disease, unlike PAH, which typically lacks overt coronary artery involvement. These findings suggest that UCP2 levels are specifically associated with dRV in PAH, but not sPHT, although it is possible that a similar role for UCP2 may be present in sPHT cases with severe rise in mPAP and perhaps significant systemic inflammation, as can be seen in obese patients with metabolic syndrome and HEpEF or HEFrEF. These data also suggest that the presence of true or functional ischemia in cRV may not be as important for the progression to dRV as currently thought.

## Discussion

Our work challenges the concept that RV failure in PAH is solely and predictably due to the increase in its afterload (i.e., mPAP) in PAH. Instead, we showed that genetic (i.e., loss-of-function UCP2 SNPs) and extra-myocardial factors (i.e., inflammation), that lead to a loss of cardiac fibroblast UCP2 levels promoting differentiation to cMFB, are associated with transition to dRV in both rodents and patients with PAH. This suggests that perhaps the focus for the treatment of PAH, that currently is restricted to targeting pulmonary vascular tone and remodeling, should expand and include treatments that may prevent or even reverse RV cMFB activation. Importantly, we suggest that the presence of UCP2 SNPs may identify patients with a predisposition to RV decompensation early after PAH diagnosis, currently not possible, facilitating optimal clinical management. Although this needs to be prospectively confirmed in multicenter cohorts, we believe our work opens a new window to much needed precision health approaches to PAH, a relatively common disease (in contrast to the rate idiopathic PAH) that remains deadly despite very expensive treatment options.

### Our animal model and mechanistic studies

A monocrotaline injection leads to severe pulmonary hypertension with an inflammatory phenotype in several organs and, following a short phase of cRV hypertrophy, causes dRV and death in ∼5-6 weeks. A complete right heart catheterization with an advancement of a Millar catheter to the main pulmonary artery to measure mPAP (as in humans), as well as simultaneous echocardiography allowed us to define hemodynamic and structural criteria (which included, among several others CO and TAPSE) to define the compensatory RV hypertrophy, in which CO and TAPSE are preserved, versus the decompensated RV, where in addition to a drop in CO and TAPSE, there is a decrease in RV systolic pressure, an increase in RA pressure and an increase in RVEDD. Thus, all monocrotaline-treated rat RV tissues were characterized as cRV or dRV, seldom done in the field where there is often no distinction between the two. This allowed us not only to observe a sharp increase in the number of cMFB but also an increase in PRMT1 levels and MICU1 methylation in both the cRV and dRV compared to control hearts; but also, that it was an ongoing decrease in UCP2 levels that specifically separated and defined the transition from cRV to dRV.

Mechanistically, we found that a central driver of fibroblast differentiation is the reduction in mCa^++^ levels, driven by the loss of UCP2 in the setting of the MICU1 hypermethylation (**Fig. 8**). Decreased mCa^++^ levels supress the activity of Ca^++^-sensitive mitochondrial enzymes (as we confirmed with PDH) and the production of diffusible metabolites that regulate the epigenetic changes involved in the change of cardiac fibroblast cell identity. Our work showed that dRV cMFB have decreased citrate and fumarate which are involved in histone acetylation via the production of acetyl-coA in the nucleus by the ATP-citrate ligase^36^. In addition, we have previously shown, translocation of mitochondrial PDH into the nucleus, directly affects histone acetylation via production of acetyl-CoA in the nucleus^37^. The PDH inhibition is in keeping with work showing that myofibroblast activation requires histone deacetylation-mediated gene repression^38,39^. To confirm the direct primary role of cMFB UCP2 loss on mCa^++^ regulation independent of MICU1 methylation, we showed that cFB from homozygote or heterozygote UCP2 KO mice have a gene-dose-dependent decreased mCa^++^. Moreover, we showed that UCP2 expression levels decreased in response to both TGF-β and TNF-α in an additive manner. Disturbances in the TGF-β axis are common in PAH, in both familiar PAH due to BMPR mutations and sporadic cases not related to known mutations^30–35^; and inflammation and cytokines like TNF-α are upregulated both in rodent and human PAH^40^. Interestingly, it has been shown by others that loss of UCP2 is associated with activation of tissue macrophages^41^ and in our UCP2 KO mice lungs we observed a significant increase in the lung inflammatory cells infiltration (also a feature of human PAH^20^). Thus, a germline UCP2 loss-of-function SNAP could affect UCP2 levels in both the bone marrow and inflammatory cells, the lungs, as well as the RV cardiac fibroblasts, producing a mutually reinforcing feedback loop between cMFB that are pro-inflammatory and circulating inflammatory cells to potentiate cMFB activation and acceleration of dRV.

**Figure 8.**
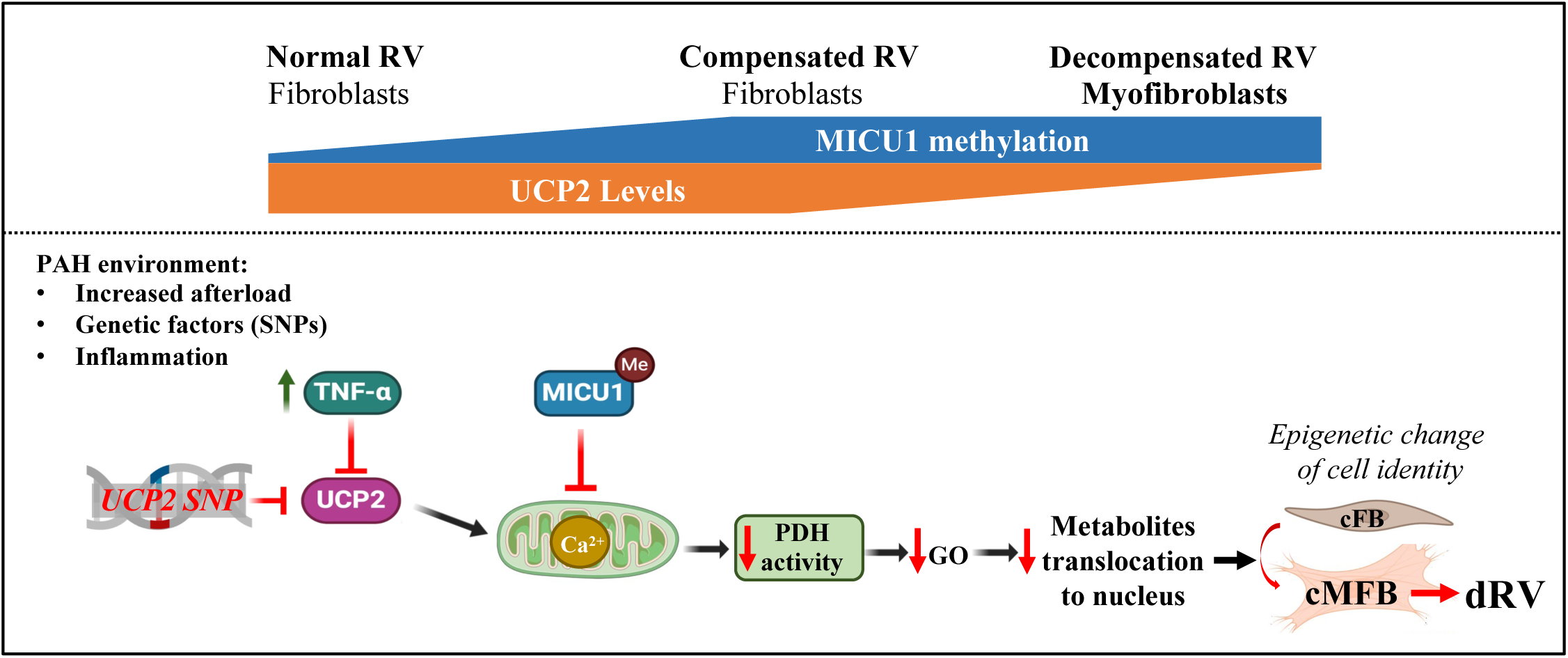
Proposed mechanism of myofibroblast activation and its contribution to right ventricular (RV) decompensation in PAH. In PAH the right ventricle (RV) undergoes afterload induced remodeling and inflammation, leading to mitochondrial dysfunction in cardiac fibroblasts (cFB). Specifically, increased methylation of MICU1 (mediated by PRMT1) and suppression of UCP2 expression by tumor necrosis factor-alpha (TNF-α) —exacerbated by UCP2 loss-of-function SNPs— reduce mitochondrial calcium uptake. This reduction inhibits pyruvate dehydrogenase (PDH) activity and impairs glucose oxidation, limiting the production of metabolites required for nuclear epigenetic signaling. These metabolic disruptions facilitate the transition from cFB to cardiac myofibroblasts (cMFB), thereby contributing to RV decompensation (as indicated by red arrows). Notably, MICU1 methylation increases primarily during the transition from control to compensated RV (cRV), whereas UCP2 downregulation is more prominent in the progression from cRV to decompensated RV (dRV). **RV,** right ventricle; **PAH,** pulmonary arterial hypertension; **SNP,** Single Nucleotide Polymorphism; **MICU1**, Mitochondrial Calcium Uptake 1; **UCP2**, uncoupling protein 2; **PDH**, pyruvate dehydrogenase; **GO**, glucose oxidation; **cFB,** cardiac fibroblasts; **cMFB,** cardiac myofibroblasts; **dRV**, decompensated right ventricle.

### Our human cohorts

included cohort 1 (Laval University): RV tissues and clinical data including cardiac index and/or TAPSE for the vast majority of patients who had been formally diagnosed with group 1 PAH (mPAP> 20 mmHg, PAWP<13 mmHg, normal left ventricular function, no thromboembolic lung or parenchymal lung disease). Cohort 2 (University of Alberta) included blood and clinical data of formally diagnosed group I PAH patients who had RHC and ECHO (performed within 2 days from each other), one year after the initiation of oral PAH therapy (PDE5 inhibitors and endothelin receptor antagonists). Cohort 3 (Duke University) included RV tissues and clinical data from heart transplant recipients with sPHT (mPAP>20 mmHg) due to ischemic or non-ischemic cardiomyopathy. We were able to detect the presence of the UCP2 SNP (rs659366) in either RV tissues or blood samples from a total of 81 patients. In these patients, we also measured RV UCP2 mRNA and protein levels as well as RV PRMT1 levels, and correlate them with CO and TAPSE. First, the negative correlation between the presence of a homozygous or heterozygous SNP with UCP2 levels confirmed that this germline SNP reflects a decrease in RV tissue UCP2 levels across all cohorts. Second, the negative correlation between UCP2 SNP and RV function (cardiac index, TAPSE) in PAH but not in sPHT patients, suggests that the UCP2 SNP is associated more with dRV than cRV in PAH and this is true even for patients that have similar mPAP. We speculate that the specificity of UCP2 levels for PAH may be due to an increase in inflammatory cytokines in PAH (particularly in patients with scleroderma), combined with higher levels of mPAP; and the fact that in patients with cardiomyopathy the RV may be affected by additional genetic factors or by ischemia.

Along with our animal data, the human studies suggest that UCP2 SNPs can be used as a biomarker of predisposition to early RV decompensation and provide some answers on why some patients with PAH decompensate faster than others independently of the rise in mPAP, based on the role of UCP2 loss in the accelerated differentiation of RV fibroblasts to myofibroblasts.

### Clinical implications

Our work suggests that biomarker-based (i.e., UCP2 SNP), precision health approaches could be applied in PAH patients, early after the establishment of the diagnosis, when it comes to anticipating, preventing or potentially reversing dRV in the future. A patient that carries the UCP2 SNP (particularly in a homozygote manner) can be followed more closely, with intense PAH and supportive therapy. Such patients should perhaps have a lower threshold for transplantation. Clinical trials with precision health design may assess whether such patients may benefit from anti-inflammatory strategies (e.g., anti-TNFα) more than patients without the SNP, when it comes to RV decompensation. At this point, anti-inflammatory strategies are only attempted with a goal to improve pulmonary vascular remodeling. The same is true with therapies targeting the TGF-β/BMPR axis (like sotatercept)^32,33^ whose targets may include RV fibroblasts as much or even more than pulmonary artery vascular cells. Our work suggests that both of these classes of therapeutics could also be applied with a goal of preventing or reversing RV decompensation. However, in such studies it may be difficult to separate vascular from cardiac effects, unless specifically designed and interpreted with this in mind. This concept may also be studied with future animal studies, using fixed RV afterload (i.e., pulmonary artery banding) models.

At this point, there is no RV-specific therapy for dRV in PAH, with the exception of perhaps mild RV inotropic effects for PDE5 inhibitors^42,43^. It is interesting that our work suggests that the answer to RV failure may not be increased RV inotropy, a reminder of the challenges of inotropes in the treatment of LV failure. Instead, we suggest that perhaps better targets may be the mitochondria (i.e., metabolic modulators like dichloroacetate^44,45^ and/or epigenetic pathways that regulate RV fibroblast differentiation^46^, along with anti-inflammatory therapies.

Finally, we found that, unexpectedly, the dRV cardiomyocytes had increased mitochondrial respiration in contrast to the dRV cMFB that had decreased oxidative phosphorylation and increased glycolysis and lactate levels. In fact, using labeled lactate we found that the CMs can use the lactate produced by neighboring cMFB as fuel to perhaps compensate to the potentially decreased ATP production. This suggests that the interpretation of functional RV metabolic imaging (for example by FDG-PET^47^), needs to be interpreted carefully and differently in the clinic, considering that the majority of the signal may come from non-cardiomyocyte cells and may not directly reflect the energetic or metabolic status of the myocardial cells as it may do in normal RVs.

### Limitations

**1.** A mouse model with germline expression of a loss-of function gene variant like the UCP2 rs659366, would be ideally needed to study the behaviour of the RV under increased afterload, in either PAH or PA banding models. Alternatively, a conditional cardiac fibroblast specific UCP2 KO could reveal a direct causal role of UCP2 loss to RV decompensation. However, in this first implication of UCP2 in RV failure, we provided data that a) show that both UCP2 mRNA and protein levels decrease in human RVs in a gene dose-dependent manner by the presence of a loss-of-function SNP; b) that TNF-α and TGF-β, both implicated in rodent and human PAH, have additive effects decreasing RV UCP2 levels; c) in a mouse UCP2 KO model (which we previously showed is associated with pulmonary hypertension) we found that RV cMFB have decreased mCa^++^ levels, the basis of the cMFB activation shown in our and in other studies (at least on the left ventricle). We believe that although our study did not prove a causal role of UCP2 loss in RV failure, enough to identify it as a therapeutic target, provides enough support to identify it as a much-needed biomarker predicting RV failure in humans, with an easy test (e.g., blood or buccal mucosa detection of SNPs with a simple PCR).

**2.** Our provocative finding that in rat RV failure the contractility of isolated cardiomyocytes is not decreased as commonly assumed, maybe biased by the technique of in-vitro sarcomere length measurement which is selecting mostly morphologically normal and beating CMs, rather than less healthy looking ones. In addition, our data were not supported by similar studies in human cardiomyocytes isolated from failing RVs, a very challenging task. We hope that our findings will inspire such important future studies in centers that can accomplish this. Nevertheless, our study is a reminder that perhaps targeting cardiomyocyte inotropy may not necessarily be an attractive option in heart failure, as shown by the many failed attempts to do this in LV failure.

## Methods

### Human tissue samples

Cohort 1: All experimental procedures using human tissues were performed with the approval of Laval University and the Institut Universitaire de Cardiologie et de Pneumologie de Québec Biosafety and Ethics Committees. All tissues were obtained from patients who had previously given written informed consent. Cohort 2: All work with human clinical data and blood samples was performed with approval from the University of Alberta Human Research Ethics Board (HREB). Blood samples were obtained from patients who had previously given written informed consent. Cohort 3: Tissue samples used for this study were were obtained from the Duke Human Heart Repository, a Duke University Hospital System Institutional Review Board-approved tissue repository. All samples were collected with written informed consent.

### Animal studies

Our work with rat and mice was performed with permission from the University of Alberta Animal Care and Use Committee (ACUC).

### Rat PHT/RV failure model

Sprague Dawley rats were purchased from the Charles River Laboratories, Canada. Rats were intraperitoneally injected with 60mg/kg monocrotaline (Sigma, C2401) at 8 weeks age. Rats underwent right heart catheterization at either 4 or 8 weeks ro identify cRV and dRV heatys respectively. However, RVs were identified as cRV or dRV only when they fulfilled strict criteria as discussed below.

### Right heart catheterization

Rats were anesthetized with 3%–4% isoflurane and maintained with 2% during procedures. A Millar catheter (microtip, 1.4F, Millar Instruments) was advanced through the jugular vein in closed-chest animals into right atrium (RA), RV and PA by guiding with a modified plastic sheath (curved). RA pressure, RV pressure and mean PA pressure (mPAP) were recorded (Power Lab, with Chart software 5.4, ADInstruments). Echocardiogram was performed immediately prior to the catheterization. Immediately after, the hearts were isolated and RV ve LV+septum dry weight was measured.

### Echocardiography

Rats were anesthetized with isoflurane and maintained a heart rate of 300-400 beats per minutes and the Vevo 3100 High Resolution Imaging System (VisualSonics, Toronto, Canada) was used. Right ventricular free wall thickness, right ventricular end-diastolic diameter and tricuspid annular plane systolic excursion (TAPSE) were recorded in the M-Mode. TAPSE was calculated by measuring the vertical movement of the tricuspid annulus between end-diastole and end-systole in the four chamber view, reflecting the longitudinal contraction of the RV. Cardiac output (CO) was calculated after determining the pulmonary artery diameter (PAD), pulmonary artery velocity time integral (PAVTI), and heart rate (HR) using the formula: CO =7.85xPAD^2^ xPAVTIxHR/10,000. Images with heart rate <300 beats/min were excluded from the analysis.

### cRV vs dRV identification

cRVs had increased RVSP and RV hypertrophy (by echo and Fulton index criteria, compared to normal: RV free wall thickness ≥1.0 mm and Fulton index ≥ 0.35) and only mildly elevated RA pressure (<7 mmHg) and normal cardiac output measured (>40 ml/min). RVs were classified as dRV if, compared to cRV, had decreased RV systolic pressure (RVSP), higher RA pressure (>7 mmHg), increased RV end diastolic diameter (RVEDD >3.5 mm), decreased TAPSE (<2 mm) and cardiac output (<40 ml/min), in keeping with reported values in rats with PHT^26^ ^48^.

### Ucp2 KO Model

We generated a WT, UCP2 Heterozygous and UCP2 KO mice from a UCP2 heterozygous colony as we have previously described^20,21^. The KO mice had exons 3-7 replaced by a larger PGK-NEO cassette. Two sets of primers are used in combination to identify wildtype (forward primer sequence: GCG TTC TGG GTA CCA TCC TA, reverse primer sequence: GCT CTG AGC CCT TGG TGT AG) and mutant mice (forward primer sequence: CTT GGG TGG AGA GGC TAT TC, reverse primer sequence: AGG TGA GAT GAC AGG AGA TC). PCRs were carried out on a MasterCycler PCR Thermal Cycler (Eppendorf Cat # 5345) by direct amplification of the DNA from ear notching biopsies with the use of Phire Tissue Direct PCR Master Mix (ThermoFisher Scientific Cat # F170) following manufacturer’s protocol. Amplifications were carried out as follows: at 94 °C for 5 min, followed by 10 cycles at 94 °C (20 sec), 65 °C (15 sec,-0.5°C per cycle decrease), 68°C (10 sec); followed by 32 cycles at 98°C (15 sec), 60°C (15 sec), 72°C (10 sec); finishing at 72°C (2 min) and holding at 4°C.

### Cardiomyocyte and cardiac fibroblast isolation

Cardiomyocyte and cardiac fibroblast isolation and culture were performed as previously described^49^. Briefly, adult cardiomyocytes were isolated from CTRL, cRV and dRV rats by digestion with type II collagenase (Worthington Biochemical Corpora-tion, LS004177) using a modified Langendorff perfusion apparatus (Harvard Apparatus, 73-4393). The cells were separated by centrifugation at 20g for 3 min (supernatant as nonmyocytes and pellet as enriched cardiomyocytes).

### Cardiomyocyte culture

After isolation, cardiomyocytes were resuspended in a stopping buffer containing 2 mM ATP followed by centrifugation at 20g for 3 min and calcium reintroduction process by sequential resuspension in a stopping buffer containing increasing concentrations of CaCl_2_ at 100, 400, and 900 mM. Before plating, culture plates were coated with laminin (Sigma-Aldrich, 11243217001) for 2 hours at room temperature, and plating medium [minimum essential medium (MEM)] with Hanks’ salts and 2 mM glutamine (Gibco, 11575032) supplemented with 10% fetal bovine serum (FBS) (Sigma-Aldrich, F1051), 10 mM 2,3-butanedione monoxime (BDM) (Sigma-Aldrich, B0753), 2 mM ATP (Sigma-Aldrich, A6419), and 1% penicilin/streptomycin/amphotericin (PSF) (Gibco, 15240-062) was prepared and kept in an incubator at 37°C with 2% CO2. Cells were plated with plating media for 1 hour, and then the medium was changed to fresh culture medium containing MEM with Hanks’ salts and 2 mM glutamine, 1% PSF, 0.1% bovine serum albumin (Sigma-Aldrich, A7906), 10 mM BDM, and 1% insulin transferrin selenium at final concentrations of 5 μg/ml for insulin, 5 μg/ml for transferrin, and 5 ng/ml for selenium (Sigma-Aldrich, I1884).

### Cardiac (myo)fibroblast culture and treatment

Non-myocytes were washed five times by centrifugation at 1500 rpm for 3 minutes using culture medium composed of Dulbecco’s Modified Eagle Medium: Nutrient Mixture F12 (DMEM/F12, Gibco, 11330032) supplemented with 100 μM ascorbic acid (Sigma, A5960), 10% fetal bovine serum (FBS, Sigma-Aldrich, F1051)), and 1% penicillin/streptomycin/amphotericin B (PSF, Gibco, 15240-062). The cells were then seeded in the same culture medium and incubated at 37°C with 5% CO₂ to allow fibroblast adhesion for 2 hours. After adhesion, the medium was replaced with fresh culture medium. When fibroblasts exhibited a spindle-shaped morphology (approximately on day 3), the medium was replaced with DMEM/F12 supplemented with 2% FBS, 100 μM ascorbic acid, and 1% PSF. All subsequent cultures and treatments were maintained in this medium. Cardiac (myo)fibroblasts at passages 1–2 were used for all experiments.

For the TNF-α dose–response experiment, cardiac fibroblasts were treated with TNF-α (Invitrogen, RTNFAI) at 0, 10, 20, or 40 ng/ml for 48 hours. For the co-treatment experiment, cells were treated with TGF-β (10 ng/ml; Cell Signaling, 5231), either alone or in combination with TNF-α (20 ng/ml), also for 48 hours.

### Confocal Imaging

Immunofluorescence staining was performed as previously described^20,37,50^ and imaging was performed using a Zeiss LSM-710 model, equipped with an Airyscan module (Carl Zeiss). Antibodies used were alpha smooth muscle actin (α-SMA) (Abcam, ab5694), Vimentin (Abcam, ab20346), Ki67 (Abcam, ab16667), PRMT1 (Cell Signaling, 2449S), ADMA (Cell Signaling, 13522S), MICU1 (Sigma Aldrich, HPA037480) and UCP2 (Cell Signaling, 89326S). All antibodies used for immunofluorescence were diluted in 1:100 and all secondary antibodies in 1:1000.

### Mitochondrial respiration measurements

c(M)FB and CM were seeded to Seahorse V7 tissue culture plates at a density of 6 × 10^4^ cells and 4,000 cells per well respectively overnight for analysis of oxygen consumption rate (OCR) using the Seahorse XFe 24 Analyzer (Agilent Technologies, Santa Clara, CA, US). The culture medium of c(M)FB and CM was removed and replaced with bicarbonate-free Seahorse XF Base medium without phenol red (Agilent Technologies, 103575-100) supplemented with 2 mM L-glutamine (Sigma-Aldrich, 607983) and 5.5 mM (baseline) D-(+)-glucose (Sigma-Aldrich, G5767). Cells were incubated in the CO2-free incubator for 1 h before being placed in the Seahorse Analyzer. OCR was measured using 3-minute mix and 3-minute measure cycles. After three baseline cycles, sequential injections of 1 µM Oligomycin, 1 µM FCCP, and 0.5 µM Rotenone/Antimycin were performed. OCR was normalized to total protein amounts isolated from each well.

### SNP genotyping assay

Genomic DNA was extracted from RV tissue or buffy coat using FlexiGene DNA Kit (QIAGEN) following manufacturer’s instruction as previously described^20^. DNA samples were quantified with a Nanodrop Spectrophotometer (ND-8000) and normalized to a concentration of 6.5 ng/μL. Samples (50ng per each) were genotyped by TaqMan® SNP Genotyping Assays for rs659366 (UCP2), and processed and read on the Droplet Digital PCR (ddPCR) QX200 (Bio-Rad) according to the manufacturer’s protocol. Each sample was partitioned into 20,000 discrete droplets and after amplification, each droplet was analyzed individually using a two-color detection system (FAM and VIC). Amplifications were carried out as follows: at 94 °C for 10 min, followed by 40 cycles at 94 °C (30 sec), 60 °C (1 min, ramp rate: 2-3°C per second); followed by holding at 98 °C (10 min), and holding at 4 °C.

### qRT-PCR and ddPCR

mRNA isolation, qRT-PCR and ddPCR were performed as previously described^20^. qRT-PCR was used to measure relative mRNA expression levels based on cycle thresholds, while ddPCR enabled absolute quantification of mRNA copy number, offering greater sensitivity—particularly useful for detecting transcripts in human samples. For ddPCR, briefly, mRNA was isolated using Qiazol (Qiagen). 100 ng of RNA were used per reaction and 18S was used as a housekeeping gene. Each sample was partitioned into 20,000 discrete droplets and after amplification, each droplet was analyzed individually using a fluorescent detection system (FAM dye). Amplifications were carried out as follows: at 48°C for 1 h, 95°C for 10 min, followed by 40 cycles at 95°C (30 s), 60°C (1 min, ramp rate: 2°C per second). Primers were purchased from Thermo Fisher Scientific (*Ucp2* Rn01754856_m1, Mm00627599_m1, Hs01075227_m1).

### Immunoblots

Tissues collecting and immunoblotting was performed as previously described^51^. Antibodies and dilutions: PRMT1 (Cell Signaling, 2449S) 1:1000, ADMA (Cell Signaling, 13522S) 1:1000, MICU1 (Sigma Aldrich, HPA037480) 1:1000, MCU (Cell Signaling, 14997S) 1:1000, UCP2 (Cell Signaling, 89326S) 1:1000, VDAC ½ (Proteintech, 10866-1-AP) and β-Actin (Santa Cruz sc81178) 1:4,000.

### Immunoprecipitation

Co-Immunoprecipitations were done using the Dynabeads Co-IP kit (Thermofisher 14321D), Dynabeads ProteinA (Thermofisher 10002D) and Pierce IP lysis buffer (Thermofisher 87787) as previously described^52^. Briefly, cells were rinsed and scraped with ice cold PBS followed by lysis with IP buffer. 50 mg equivalent of cell pellet volume were added to 1 µg of antibody or the appropriate IgG conjugated to beads and incubated at either 4°C for 2 hrs (for Flag IPs) or overnight at 4°C (for other IPs). Beads were then washed, eluted and boiled in 4x Laemmli buffer (Biorad).

### Mitochondrial Calcium (mCa^++^)

was measured by using Rhod-2 AM. Cells were incubated with DMEM/F12 media with 5 μM Rhod-2 AM for 30min at 37°C, followed by an incubation with serum-free DMEM/F12 medium with 200 nM MitoTracker Green to detect the localization of mitochondria for another 30min. Then a confocal fluorescence microscope (Carl Zeiss) was used to evaluate the mCa^++^ levels. Rhod-2 was excited at 514 nm, and emission was monitored at 530-580 nm. MitoTracker Green was excited at 488 nm, and emission was monitored at 505-535 nm.

### Mass Spectrometry

Mass spectrometry was performed as previously described^51^. Briefly, c(M)FB and CM were cultured in triplicates in 60 mm dishes overnight. c(M)FB were treated with 17.5mM C-13 glucose in no-phenol red, no-glucose DMEM for 12 h, and CM were treated with 5mM C-13 Lactate for 2h. Then cells were washed and scraped in 1 mL of cold PBS solution, followed by centrifugation at 550 g for 5 min at 4°C. Cell pellets were re-suspended in an ice-cold mixture of methanol and ddH2O (80:20 v/v) containing internal standards (C/D/N Isotopes, Quebec, Canada). Samples were periodically vortexed, then sonicated (10 pulses, 50% intensity) and centrifuged at 13,000 rpm for 10 min at 4oC. Supernatants were collected, and the extraction procedure was repeated one more time. The combined supernatants containing metabolites were transferred to a new set of tubes and dried using a SpeedVac vacuum concentrator (Thermo Fisher Scientific). 200ul c(M)FB supernatants were collected and mixed with 800ul methanol and the following steps were the same as cell metabolites extraction. The dried metabolites were resuspended in 100 uL of LC/MS-grade water/acetonitrile (60:40 v/v) (Fisher Scientific) before Mass Spec analysis. The extracts from supernatant were diluted in water/acetonitrile (60:40, v/v) 10 times before HPLC/MS analysis. Standard and sample solutions were analyzed using an Aria MX HPLC system (Thermo Fisher Scientific) coupled to Orbitrap Elite Mass spectrometer (Thermo Fisher Scientific). The Xcalibur software v. 2.2 (Thermo Fisher Scientific) was used for data acquisition and analysis. For the determination of Lactate and TCA cycle metabolites, a ten-point calibration curve of target compounds with a concentration range from 0.005 mg/mL to 25 mg/mL was constructed based on the peak area ratio of the target compounds/internal standard vs. the target compounds concentration. The target compounds concentration was extrapolated from this calibration curve.

### PDH activity assay

PDH activity was measured using commercially available kits (Abcam, ab109882) following the manufacturer’s manual. As the protocol recommended, 50 ul blocking solution was added to an empty well of the microplate and 250ug protein in 50ul buffer were added to the same well and mixed well by pipetting. Mildly add a dipstick to the microplate to allow the sample to absorb on the dipstick for 20 min. Then, wash the well with 40 µl sample buffer for 10 min and add 300 ul of activity buffer to an empty well for each dipstick. Transferring the dipstick into the activity buffer to react for 20 min. After signal development, the dipstick was washed with deionized water for 5 min. The dipstick was visualized on a scanner and the signal was quantified using the Image J program

### Isolated RV Langendorff Perfusion (Ex-vivo RV contractility)

RV pressure of isolated hearts was measured by Langendorff perfusion system as previously described^42^. Rats were anesthetized with isoflurane. A midline sternotomy was performed, and within 1 minute, the heart was isolated and the aorta was cannulated and perfused with Krebs’ buffer (NaCl 118.5mM, NaHCO_3_ 25mM, KCl 4.7mM, MgSO_4_ 1.2mM, KH_2_PO_4_ 1.2mM, CaCl_2_ 1.4mM, Glucose 5.5mM) at 12 to 13 ml/min. The hearts had a mean intrinsic rate of 240 to 300 bpm. A pacemaker was connected to the myocardium with a pacing rate at 300/min. A 0.03 cm^3^ latex balloon (Harvard Apparatus, Saint-Laurent, Quebec, Canada) filled with water and attached to a pressure transducer (Cobe, Richmond Hill, Ontario, Canada) was placed in the RV via the right atrium, and pressure traces were sampled at a rate of 1000 Hz by PowerLab. Pressure readings were converted into first-derivative traces and were analyzed with Chart 8 software (ADInstruments Inc, Colorado Springs, Colo).

### In vitro assessment of cell contractility

Shortening of sarcomere was assessed by a video-based edge-detection system (IonOptix, Milton, MA, USA). RV myocytes were plated on a round cover glass (25mm) coated with laminin. After 2 hours incubation, the cover glass was glued to an experimental chamber with a pair of electrodes placed inside and perfused with Krebs’ buffer (same as EX-vivo) at 1 ml/min. Cells were stimulated with 10 volts at a frequency of 2 Hz (1msec duration) using a field stimulator. The relaxed sarcomere length (relaxed length, RL) and shortest sarcomere length during the contraction (contracted length, CL) were recorded. The sarcomere shortening was calculated using the formula: % shortening = (RL-CL)/RL*100%.

### Statistical analysis

All statistical analyses performed on GraphPad Prism 10 (GraphPad Software, La Jolla, US). Values are expressed as mean ± SEM. The use of parametric or non-parametric tests was decided after assessment of normal distribution of the values by the Shapiro-Wilk normality test. For parametric tests, we used two-tailed, unpaired Student’s t test to assess statistical significance between two groups, and multiple groups were compared using one-way analysis of variance (ANOVA) or two-way repeated measures ANOVA followed by a Bonferroni post hoc analysis. For non-parametric tests, Mann-Whitney U test was used for comparisons between two groups and Kruskal-Wallis test was used to compare multiple groups. Non-parametric correlation analysis of the clinical data was performed using the Spearman correlation co-efficient. Significance for all statistical testing was p < 0.05.

### Sources of Funding

Y. Zhang is supported by Paroian Family PH Scholarship. G. Sutendra is supported by an Alberta Innovates Translational Health Chair and a Canada Research Chair (tier 2), along with grants from Canadian Institutes of Health Research. E. Michelakis is supported by a Canada Research Chair, and this work was supported by the Heart and Stroke Foundation of Canada, and Canadian Institutes of Health Research.

## Disclosures

None

---

**Supplementary Table1.**
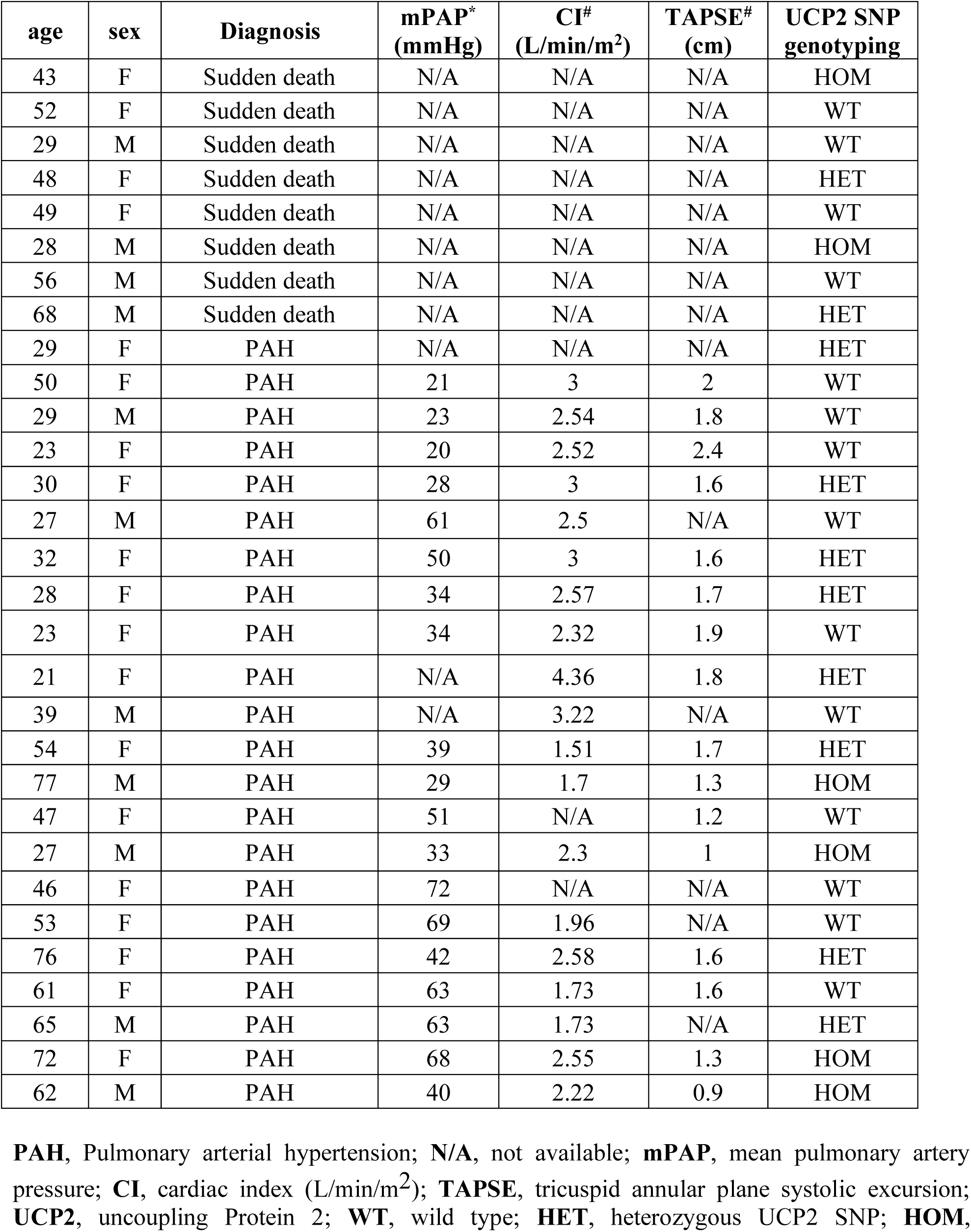
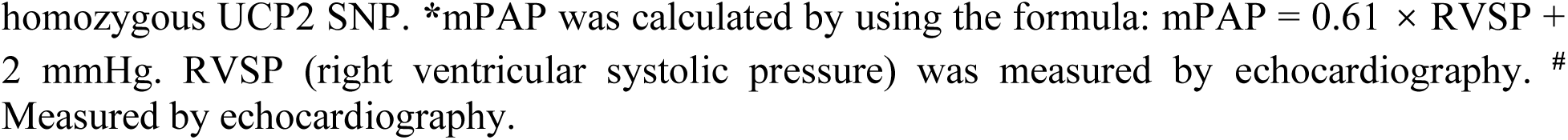
Clinical information of the first cohort of patients with pulmonary arterial hypertension (cohort 1).

| **age** | **sex** | **Diagnosis** | **mPAP\***<br>**(mmHg)** | **CI#**<br>**(L/min/m<sup>2</sup>)** | **TAPSE#**<br>**(cm)** | **UCP2 SNP**<br>**genotyping** |
| --- | --- | --- | --- | --- | --- | --- |
| 43 | F | Sudden death | N/A | N/A | N/A | HOM |
| 52 | F | Sudden death | N/A | N/A | N/A | WT |
| 29 | M | Sudden death | N/A | N/A | N/A | WT |
| 48 | F | Sudden death | N/A | N/A | N/A | HET |
| 49 | F | Sudden death | N/A | N/A | N/A | WT |
| 28 | M | Sudden death | N/A | N/A | N/A | HOM |
| 56 | M | Sudden death | N/A | N/A | N/A | WT |
| 68 | M | Sudden death | N/A | N/A | N/A | HET |
| 29 | F | PAH | N/A | N/A | N/A | HET |
| 50 | F | PAH | 21 | 3 | 2 | WT |
| 29 | M | PAH | 23 | 2.54 | 1.8 | WT |
| 23 | F | PAH | 20 | 2.52 | 2.4 | WT |
| 30 | F | PAH | 28 | 3 | 1.6 | HET |
| 27 | M | PAH | 61 | 2.5 | N/A | WT |
| 32 | F | PAH | 50 | 3 | 1.6 | HET |
| 28 | F | PAH | 34 | 2.57 | 1.7 | HET |
| 23 | F | PAH | 34 | 2.32 | 1.9 | WT |
| 21 | F | PAH | N/A | 4.36 | 1.8 | HET |
| 39 | M | PAH | N/A | 3.22 | N/A | WT |
| 54 | F | PAH | 39 | 1.51 | 1.7 | HET |
| 77 | M | PAH | 29 | 1.7 | 1.3 | HOM |
| 47 | F | PAH | 51 | N/A | 1.2 | WT |
| 27 | M | PAH | 33 | 2.3 | 1 | HOM |
| 46 | F | PAH | 72 | N/A | N/A | WT |
| 53 | F | PAH | 69 | 1.96 | N/A | WT |
| 76 | F | PAH | 42 | 2.58 | 1.6 | HET |
| 61 | F | PAH | 63 | 1.73 | 1.6 | WT |
| 65 | M | PAH | 63 | 1.73 | N/A | HET |
| 72 | F | PAH | 68 | 2.55 | 1.3 | HOM |
| 62 | M | PAH | 40 | 2.22 | 0.9 | HOM |

**PAH**, Pulmonary arterial hypertension; **N/A**, not available; **mPAP**, mean pulmonary artery pressure; **CI**, cardiac index (L/min/m<sup>2</sup>); **TAPSE**, tricuspid annular plane systolic excursion; **UCP2**, uncoupling Protein 2; **WT**, wild type; **HET**, heterozygous UCP2 SNP; **HOM**,
homozygous UCP2 SNP. \*mPAP was calculated by using the formula: $mPAP = 0.61 \times RVSP + 2$ mmHg. RVSP (right ventricular systolic pressure) was measured by echocardiography. # Measured by echocardiography.

**Supplementary Table2.** Clinical information of the second cohort of patients with pulmonary arterial hypertension (cohort 2).

| age | sex | Diagnosis | mPAP*<br>(mmHg) | CI#<br>(L/min/m<sup>2</sup>) | TAPSE#<br>(cm) | UCP2 SNP<br>genotyping |
| --- | --- | --- | --- | --- | --- | --- |
| 35 | F | PAH | 39 | 3.9 | 1.8 | WT |
| 38 | F | PAH | 44 | 3.9 | 2.5 | WT |
| 44 | M | PAH | 42 | 4 | 2.6 | WT |
| 42 | F | PAH | 46 | 4.1 | 2.7 | WT |
| 33 | F | PAH | 51 | 3.7 | 2.4 | WT |
| 49 | F | PAH | 52 | 3 | 1.7 | WT |
| 55 | M | PAH | 55 | 3.3 | 1.5 | WT |
| 45 | F | PAH | 62 | 2.8 | 1.6 | WT |
| 59 | M | PAH | 49 | 3.6 | 2.1 | WT |
| 61 | F | PAH | 54 | 3 | 1.5 | WT |
| 42 | F | PAH | 48 | 2.9 | 1.6 | HET |
| 35 | F | PAH | 61 | 2.6 | 1.2 | HET |
| 36 | F | PAH | 47 | 2.8 | 2 | HET |
| 39 | M | PAH | 55 | 2.6 | 1.8 | HET |
| 37 | F | PAH | 60 | 2.5 | 1.4 | HET |
| 39 | F | PAH | 67 | 2.6 | 1.3 | HET |
| 33 | F | PAH | 50 | 2.8 | 1.4 | HOM |
| 35 | M | PAH | 52 | 2.4 | 1.5 | HOM |
| 39 | F | PAH | 54 | 2.5 | 1.3 | HOM |
| 44 | F | PAH | 62 | 1.8 | 1 | HOM |
| 34 | F | PAH | 48 | 2 | 1.5 | HOM |

A cohort of 21 consecutive PAH patients (with normal systolic and diastolic LV function, PAWP<13mmHg) referred to the PAH clinic and treated with identical PAH therapy, had right heart catheterization and echocardiography within 48 hrs from each other, approximately 12 months after their initial PAH diagnosis.
**PAH**, Pulmonary arterial hypertension; **mPAP**, mean pulmonary artery pressure; **CI**, cardiac index (L/min/m<sup>2</sup>); **TAPSE**, tricuspid annular plane systolic excursion; **UCP2**, uncoupling Protein 2; **WT**, wild type; **HET**, heterozygous UCP2 SNP; **HOM**, homozygous UCP2 SNP. \* mPAP was measured by closed-chest right heart catheterization. #Measured by echocardiography.

**Supplementary Table3.** Clinical information of a cohort of patients with secondary pulmonary hypertension (cohort 3).

| age | sex | Diagnosis | mPAP*<br>(mmHg) | CI#<br>(L/min/m<sup>2</sup>) | TAPSE#<br>(cm) | UCP2 SNP<br>genotyping |
| --- | --- | --- | --- | --- | --- | --- |
| 51 | M | NICM | 14 | 3.7 | 1.1 | HET |
| 69 | M | ICM | 19 | 2.5 | 1.4 | WT |
| 64 | M | NICM | 18 | 2.8 | 1.3 | HET |
| 55 | M | ICM | 16 | 2.2 | 1.1 | WT |
| 56 | F | ICM | N/A | N/A | N/A | WT |
| 55 | M | NICM | 19 | 2.7 | N/A | HET |
| 62 | M | NICM | 17 | 2.6 | N/A | HET |
| 61 | M | HCM | 14 | 1.7 | 1.2 | WT |
| 57 | M | NICM | N/A | N/A | N/A | WT |
| 69 | M | ICM | 27 | 2.4 | N/A | WT |
| 63 | M | NICM | 46 | 2.8 | 1.5 | WT |
| 45 | M | NICM | 39 | 3 | N/A | HET |
| 65 | M | ICM | 39 | 2.5 | N/A | WT |
| 34 | F | NICM | 37 | 2.6 | 1.7 | HET |
| 66 | M | ICM | 40 | 2.1 | 1.5 | HET |
| 61 | M | NICM | 30 | 2.8 | 2.7 | WT |
| 63 | F | NICM | 21 | 2.3 | 2.9 | HET |
| 42 | M | NICM | 20 | 2.4 | N/A | HOM |
| 50 | M | NICM | 40 | 1.5 | 1.8 | HOM |
| 55 | M | NICM | 24 | 1.9 | 1.9 | WT |
| 63 | M | NICM | 22 | 1.5 | 1.1 | WT |
| 61 | F | ICM | 21 | 2.6 | 1.36 | WT |
| 47 | M | NICM | 24 | 2.1 | 0.4 | HET |
| 66 | F | NICM | 25 | 1.4 | 1.3 | HET |
| 68 | M | ICM | 40 | 2.5 | 1.4 | HOM |
| 63 | M | NICM | 39 | 1.9 | 1.4 | HOM |
| 56 | M | NICM | 40 | 1.5 | 1.3 | WT |
| 58 | M | ICM | 29 | 2.1 | 0.6 | HET |
| 69 | M | ICM | 42 | 1.7 | 1.1 | HET |
| 51 | M | NICM | 25 | 2.11 | 1.3 | HET |

**PHT**, pulmonary hypertension; **N/A**, not available; **CI**, cardiac index (L/min/m<sup>2</sup>); **TAPSE**, tricuspid annular plane systolic excursion; **UCP2**, uncoupling Protein 2; **NICM**, Non-ischemic Cardiomyopathy; **ICM**, Ischemic Cardiomyopathy; **WT**, wild type; **HET**, heterozygous UCP2
SNP; **HOM**, homozygous UCP2 SNP. \*Measured by closed-chest right heart catheterization.

**Supplementary figure 1.**
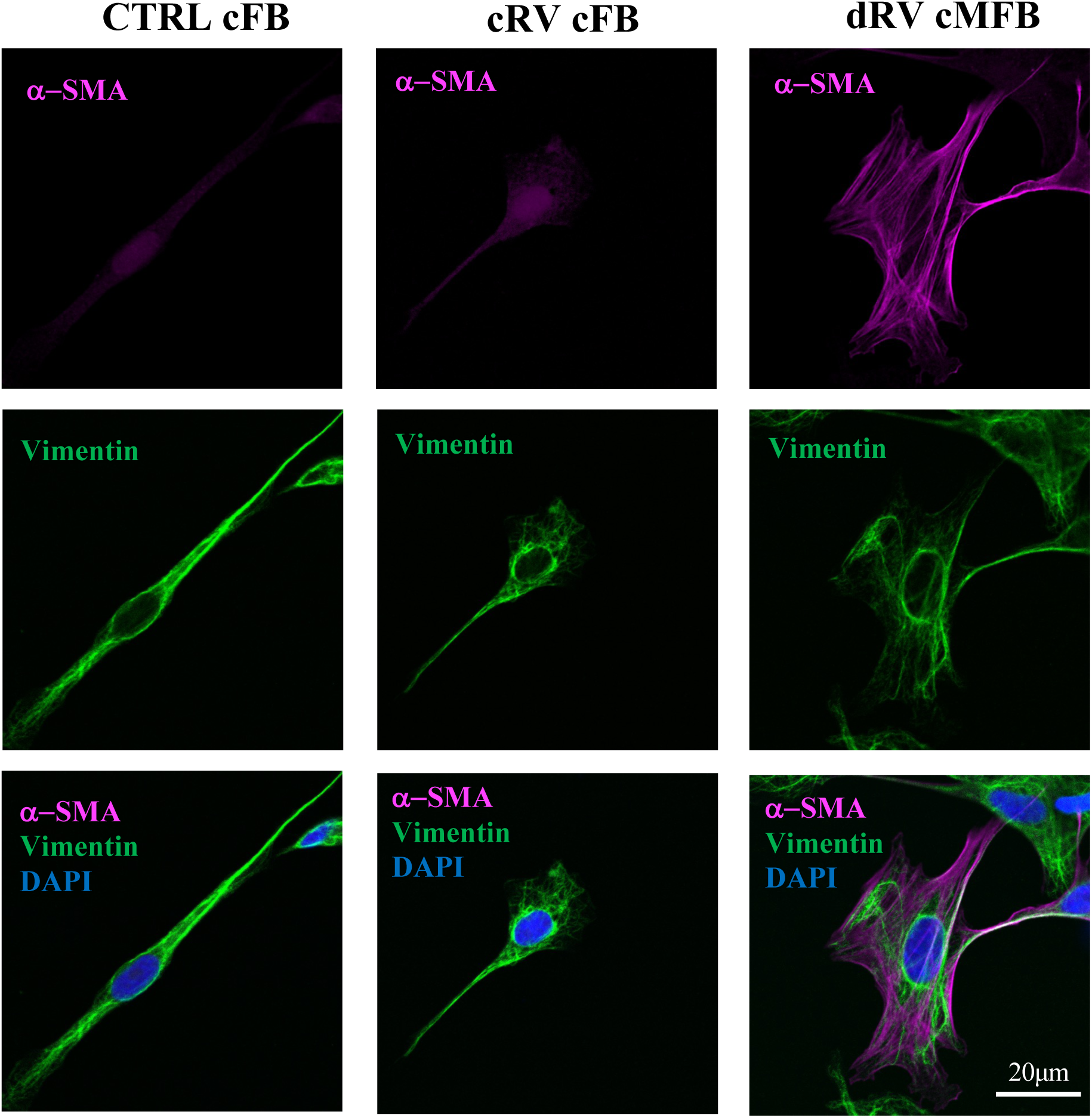
Cardiac (myo)fibroblasts from decompensated RV have elevated *α*-SMA expression and a very different cell shape relative to control and compensated RV (Confocal microscopy) **CTRL**, control (normal right ventricle, without monocrotaline injection); **cRV**, compensated right ventricle; **dRV**, decompensated right ventricle; **cFB**, cardiac fibroblasts; **cMFB**, cardiac myofibroblasts; **α-SMA**, alpha smooth muscle actin; **DAPI**, 4′,6-diamidino-2-phenylindole (labels the nucleus).

**Supplementary figure 2.**
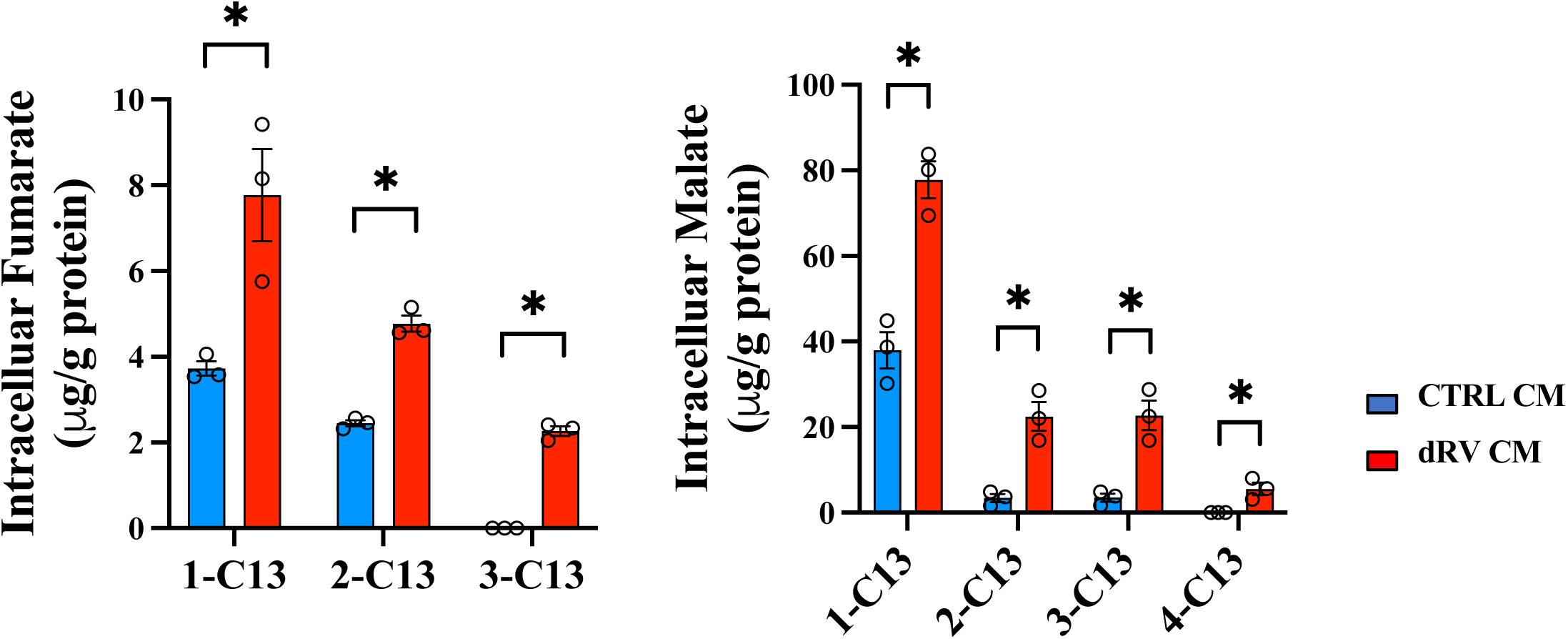
Enhanced lactate utilization by cardiomyocytes in decompensated right ventricles. Mass Spectrometry shows the intracellular labeled fumarate and malate levels in cardiomyocytes from normal RV and dRV after treatment with C-13-lactate (5mM) for 12h (1-C13, 2-C13, 3-C13 and 4-C13 signifies the number of carbons labelled by C13). Values are expressed as mean ± SEM; n = 3 animals for each group. *P < 0.05. These comparisons was made using the Mann-Whitney U test. **CTRL**, control (normal right ventricle, without monocrotaline injection); **dRV**, decompensated right ventricle; **CM**, cardiomyocytes.

**Supplementary figure 3.**
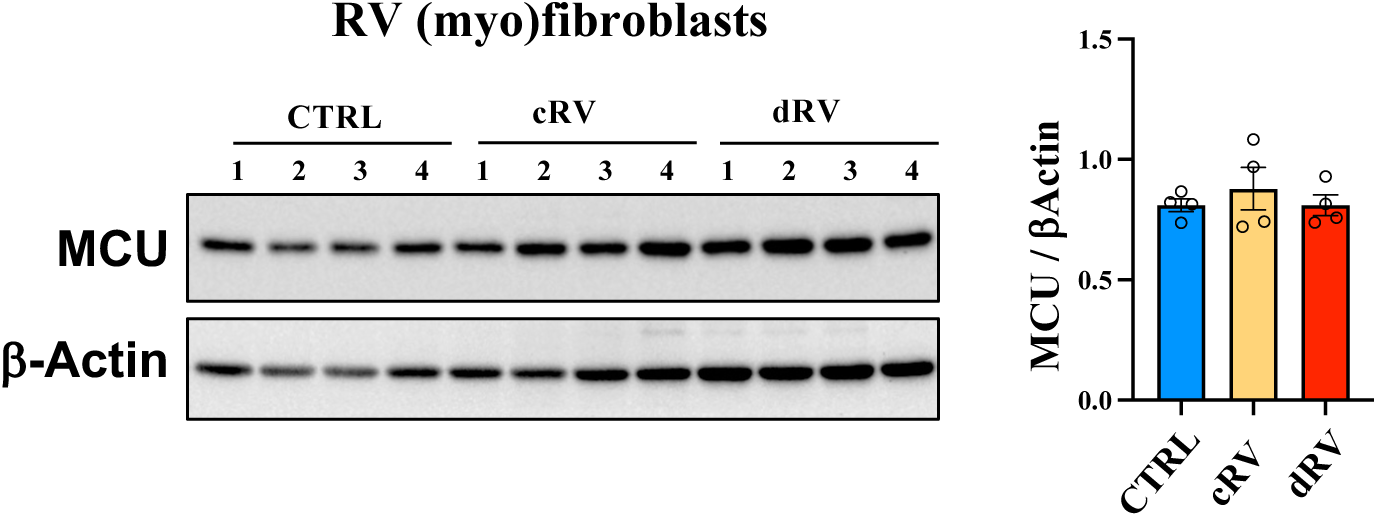
MCU levels are not different among (myo)fibroblasts from control, cRV, and dRV. Immunoblot shows the amount of MCU and Β-Actin in (myo)fibroblasts from normal RV, cRV and dRVs. Quantification values are expressed as mean ± SEM; n = 4 animals for each group. The comparison was made using Kruskal–Wallis tests. **CTRL**, control (normal right ventricle, without monocrotaline injection); **cRV**, compensated right ventricle; **dRV**, decompensated right ventricle; **MCU**, mitochondrial calcium uniporter.

**Supplementary figure 4.**
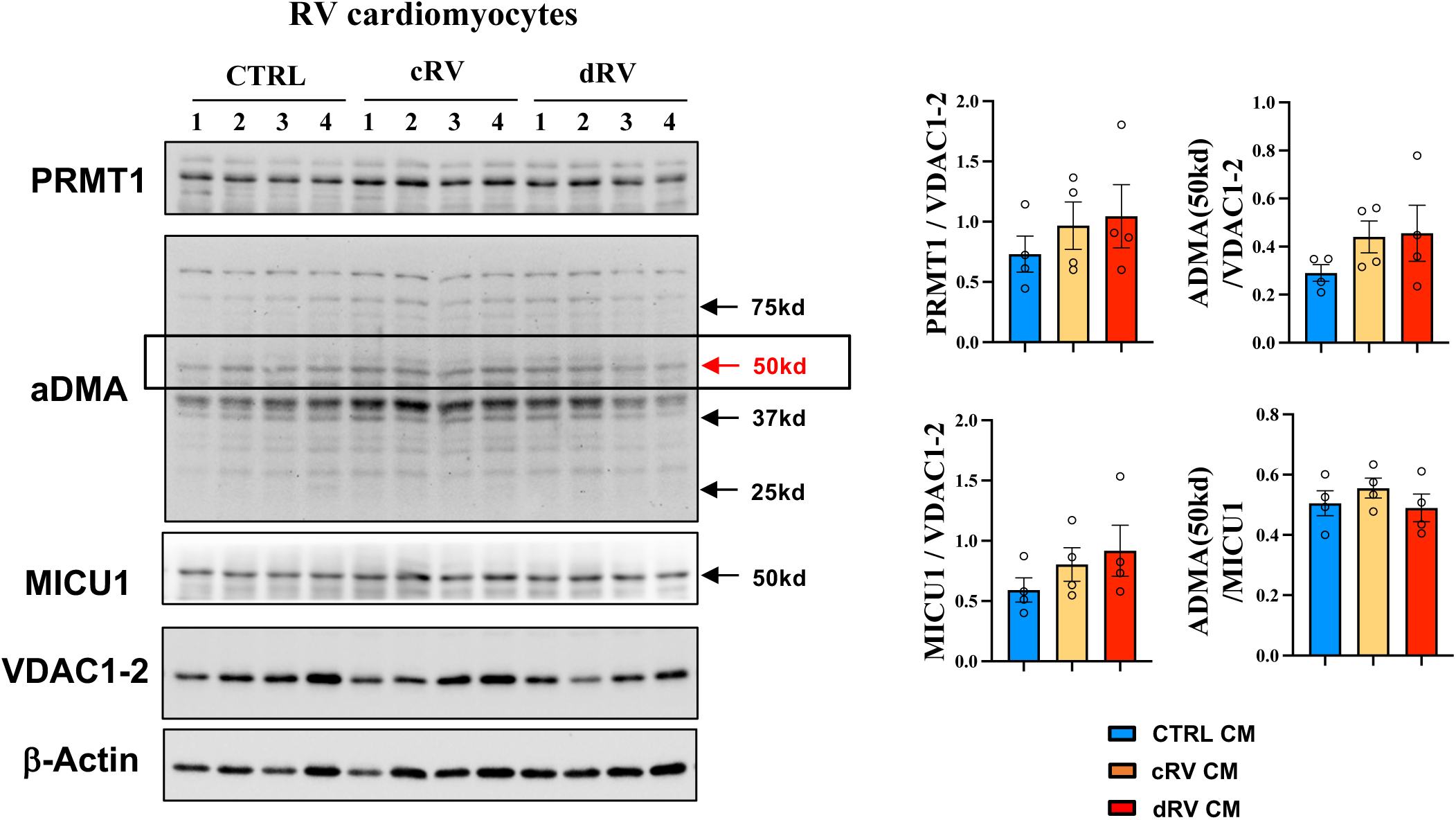
The status of cardiomyocyte MICU1 methylation does not change in (myo)fibroblasts from control to cRV to dRV. Immunoblot shows the amount of PRMT1, ADMA, MICU1, VDAC1/2 and β-Actin in cardiomyocytes from normal RV, cRV and dRVs. Quantification values are expressed as mean ± SEM; n = 4 animals for each group. Comparisons were made using Kruskal–Wallis tests. **CTRL**, control (normal right ventricle, without monocrotaline injection); **cRV**, compensated right ventricle; **dRV**, decompensated right ventricle; **PRMT1**, Protein Arginine Methyltransferase 1; **ADMA**, Asymmetric dimethylarginine; **MICU1**, Mitochondrial Calcium Uptake 1; **VDAC**, Voltage-Dependent Anion Channels; **CM**, cardiomyocytes.

**Supplementary Figure 5.**
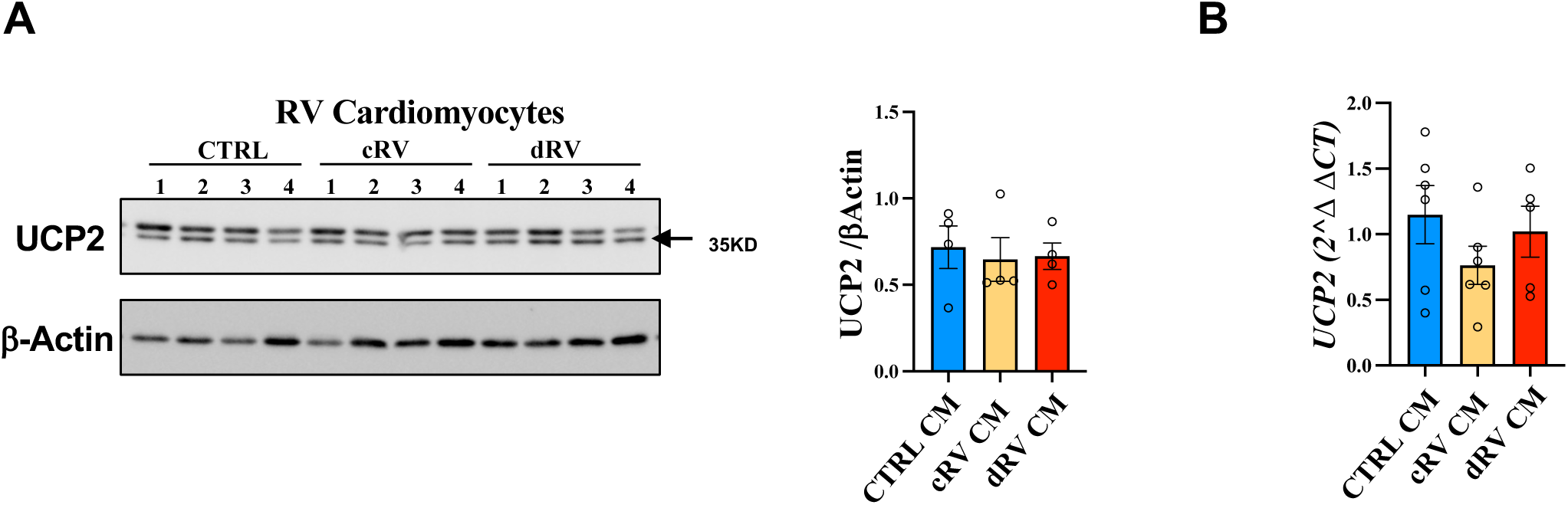
UCP2 levels remain unchanged in cardiomyocytes from control to cRV to dRV. **A.** Immunoblot (left) shows the UCP2 protein level in right ventricular cardiomyocytes from normal RV, cRV and dRV from PHT rats. Quantification values (right) normalized by β-Actin are expressed as mean ± SEM; n = 5 animals for each group. **B.** q-PCR shows the UCP2 RNA levels in right ventricular cardiomyocytes from normal RV, cRV and dRV from PHT rats. These comparisons were made using one-way Kruskal–Wallis tests. **CTRL**, control (normal right ventricle, without monocrotaline injection); **cRV**, compensated right ventricle; **dRV**, decompensated right ventricle; **UCP2**, uncoupling Protein 2; **CM**, cardiomyocytes.

**Supplementary Figure 6.**
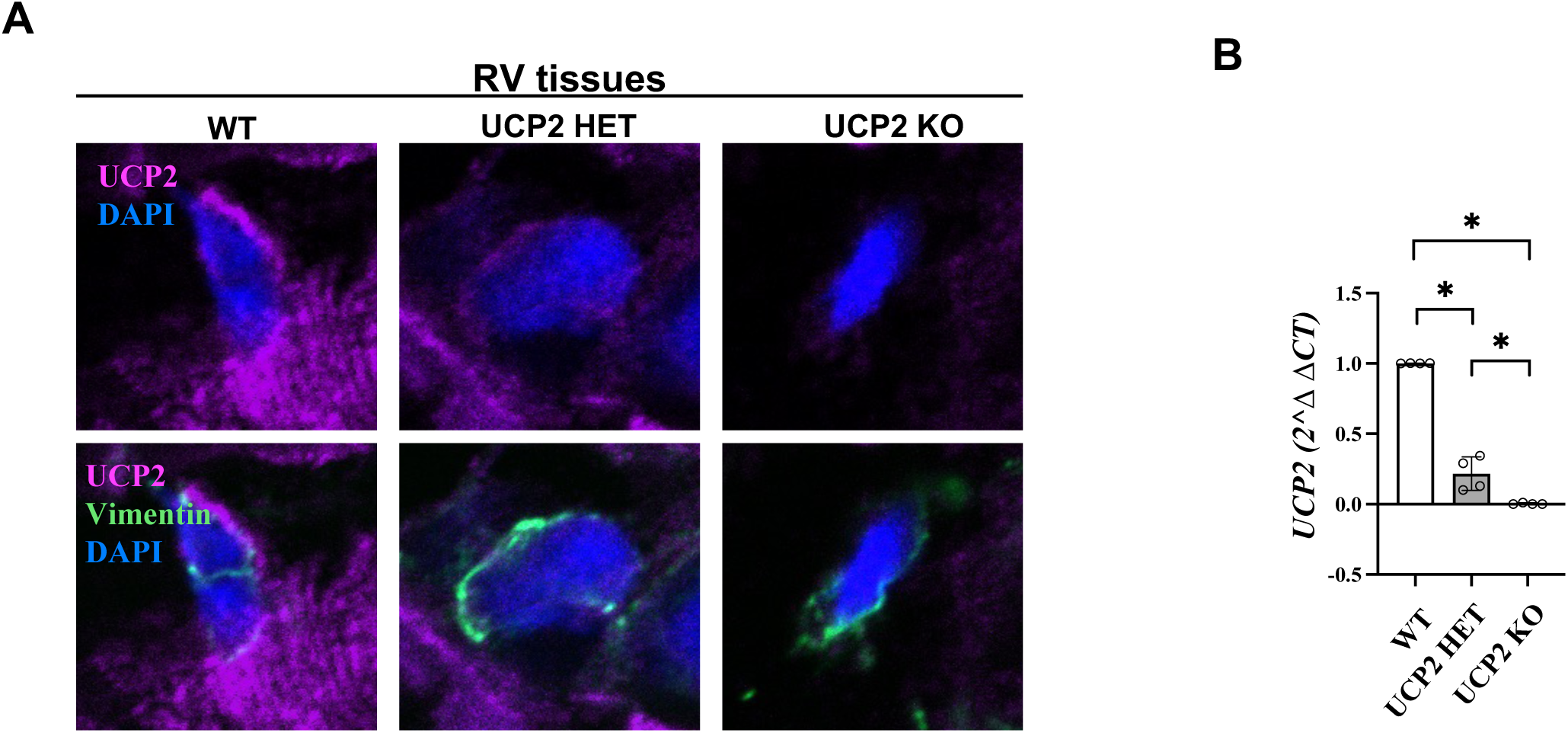
UCP2 knockout (myo)fibroblasts show no detectable UCP2 protein or RNA expression. **A.** Representative confocal images show the UCP2 immunofluorescence intensity in RV cFBs from WT, UCP2 HET and UCP2 KO mice, confirming the genotype for the Ucp2 mutant mice. **B.** qRT-PCR (from heart tissues) shows that the UCP2 knockout mice have no detectable levels of Ucp2 mRNA, compared to wild type controls, while the UCP2 heterozygous mice express ∼30% mRNA levels. This comparison was made using Kruskal–Wallis tests followed by pairwise Mann–Whitney U tests. **CTRL**, control (normal right ventricle, without monocrotaline injection); **cRV**, compensated right ventricle; **dRV**, decompensated right ventricle; **UCP2**, uncoupling Protein 2; **DAPI**, 4′,6-diamidino-2-phenylindole (labels the nucleus); **WT,** wild type; **HET,** heterozygous; **KO**, Knockout.

**Supplementary Figure 7.**
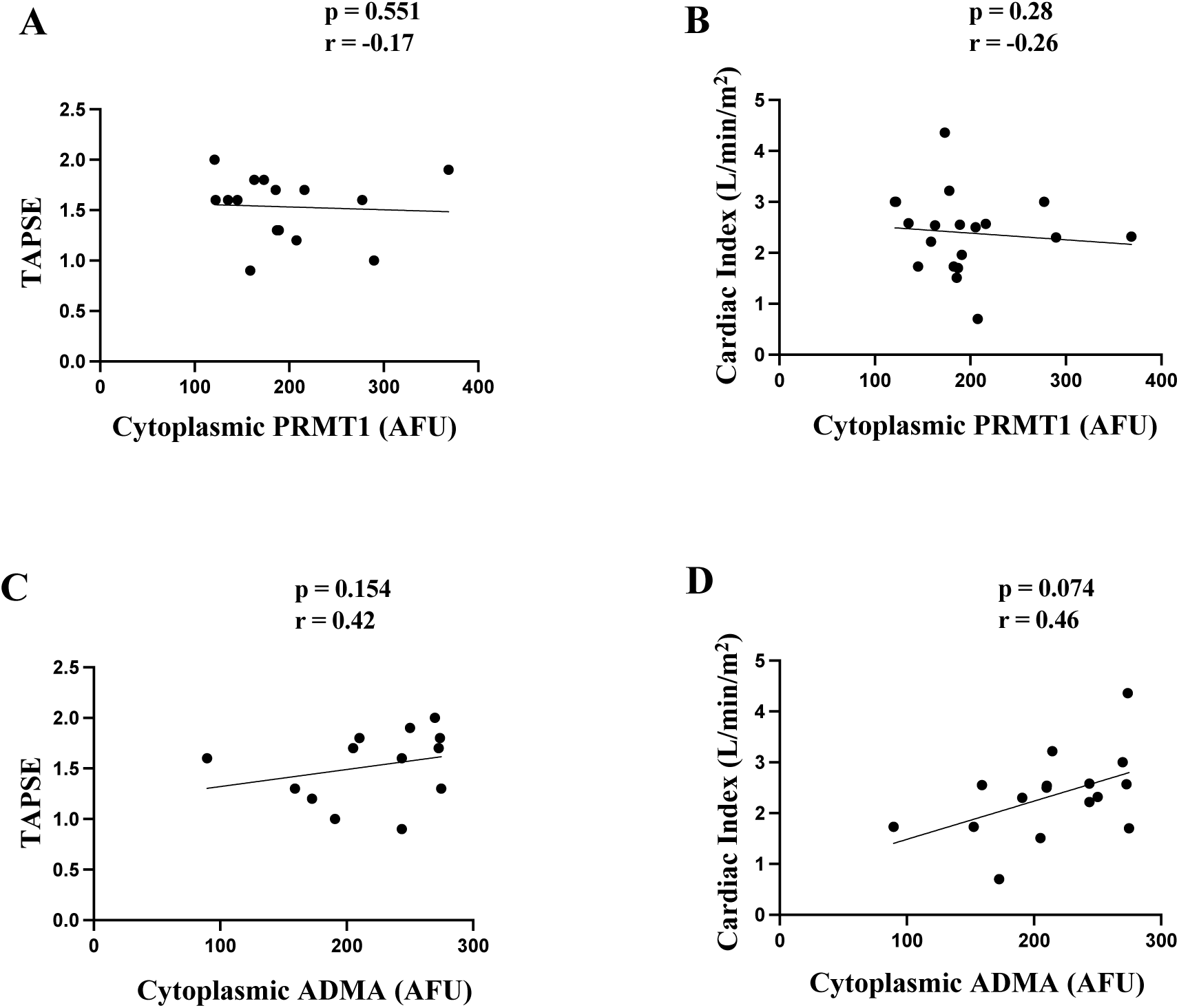
Arginine methylation status does not correlate with TAPSE or cardiac index in patients with PAH. **A.** Spearman correlation test shows the correlation between cytoplasmic PRMT1 immunofluorescence intensity in (myo)fibroblasts with TAPSE in cohort 1 patients with PAH. Scatter dot plots represent individual values. *p<0.05 **B.** Spearman correlation test shows the correlation between cytoplasmic PRMT1 immunofluorescence intensity in (myo)fibroblasts with Cardiac Index in cohort 1 patients with PAH. Scatter dot plots represent individual values. **p<0.01 **C.** Spearman correlation test shows the correlation between cytoplasmic ADMA immunofluorescence intensity in (myo)fibroblasts with TAPSE in cohort 1 patients with PAH. Scatter dot plots represent individual values. *p<0.05 **D.** Spearman correlation test shows the correlation between cytoplasmic ADMA immunofluorescence intensity in (myo)fibroblasts with Cardiac Index in cohort 1 patients with PAH. Scatter dot plots represent individual values. **p<0.01 **PRMT1**, Protein Arginine Methyltransferase 1; **ADMA**, Asymmetric dimethylarginine. **TAPSE,** Tricuspid Annular Plane Systolic Excursion.

**Supplementary Figure 9.**
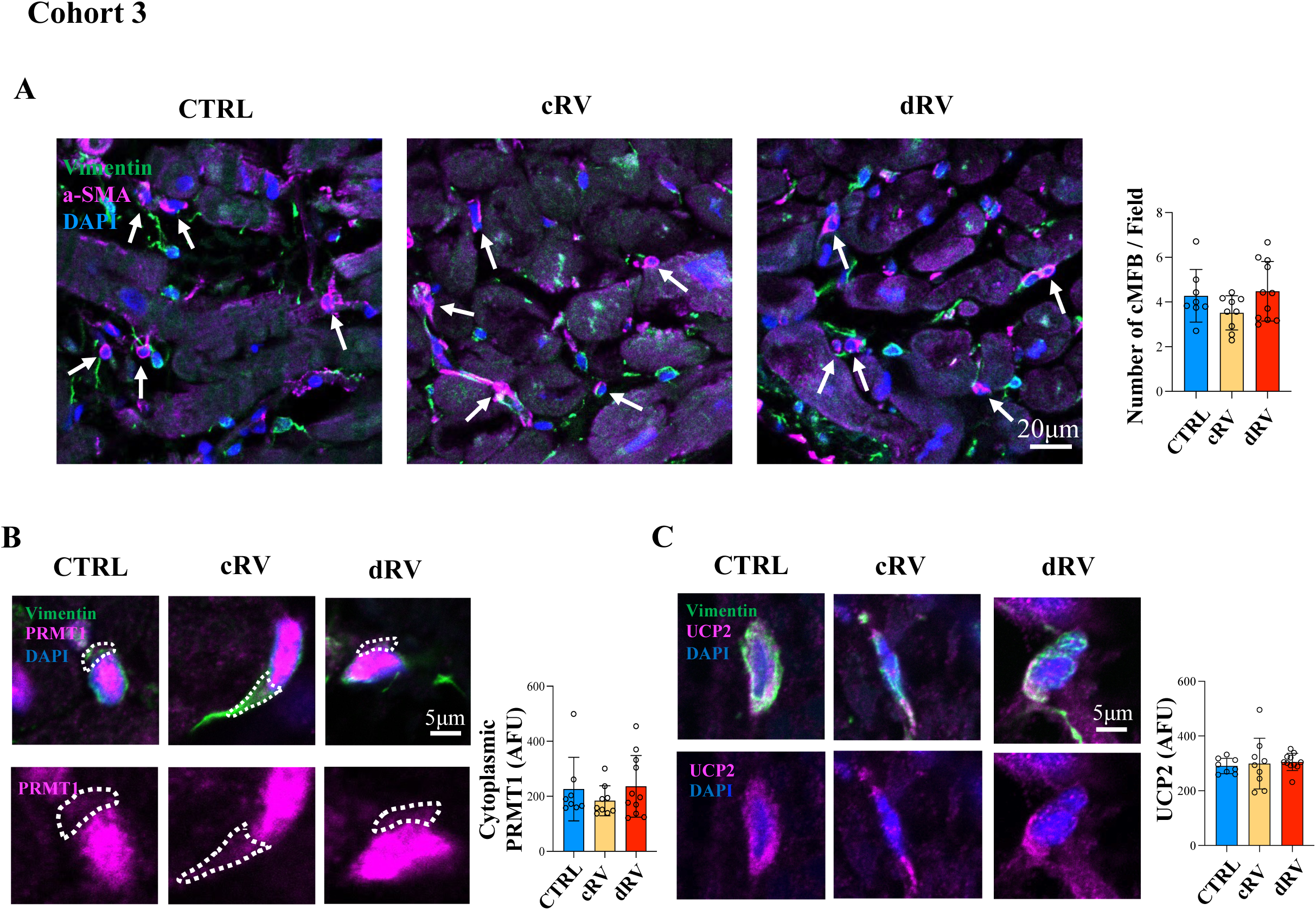
The number of myofibroblasts, cytoplasmic PRMT1, and UCP2 levels in fibroblasts are not different among control, cRV, and dRV samples from patients with secondary pulmonary hypertension (Cohort 3). **A.** Confocal microscopy (left) shows the number of myofibroblasts (positive with both α-SMA and Vimentin) from paraffin-embedded right ventricle sections from CTRL subjects (n=8), cRV (n=10) and dRV (n=11) patients with secondary pulmonary hypertension (sPHT). Scatter dot plots represent individual values. Quantification values (**right**) are expressed as mean ± SEM. **B.** Confocal microscopy (left) shows the cytoplasmic PRMT1 level in (myo)fibroblasts (vimentin-positive cells) from paraffin-embedded right ventricle sections from CTRL subjects (n=8), cRV (n=10) and dRV (n=11) cohort 3 patients with secondary pulmonary hypertension (sPHT). The dotted line tracing follows the Vimentin signal in order to trace the cytoplasm. Scatter dot plots represent individual values. Quantification values (**right**) are expressed as mean ± SEM. **C.** Confocal microscopy (left) shows the UCP2 level in (myo)fibroblasts (vimentin-positive cells) from paraffin-embedded right ventricle sections from CTRL subjects (n=8), cRV (n=10) and dRV (n=11) cohort 3 patients with secondary pulmonary hypertension (sPHT). Scatter dot plots represent individual values. Quantification values (**right**) are expressed as mean ± SEM. These comparisons were made using Kruskal–Wallis tests. **CTRL**, Normal right ventricle but with left ventricular failure, **cRV**, compensated right ventricle; **dRV**, decompensated right ventricle; **α-SMA**, alpha smooth muscle actin; **DAPI**, 4′,6-diamidino-2-phenylindole (labels the nucleus); **PRMT1**, Protein Arginine Methyltransferase 1; **UCP2,** uncoupling protein 2.

## Notes

### Competing Interest Statement

The authors have declared no competing interest.

